# A Two-Phase Core-Plasma Model for Microvascular Blood Flow: Comparative Analysis of Hemodynamic Models

**DOI:** 10.1101/2025.06.25.661657

**Authors:** Maya Salame, Marianne Fenech

**Affiliations:** Department of Mechanical Engineering, University of Ottawa, Ottawa, Ontario, Canada

## Abstract

Microcirculatory blood flow exhibits complex non-Newtonian behavior, including shear-thinning properties and the formation of a cell-free layer (CFL)—a plasma-rich region near vessel walls. While traditional rheological models such as Newtonian, Power Law, and Carreau describe certain flow characteristics, and empirical models like the double-parameter power fit have been used to capture velocity profiles, these approaches fall short in fully characterizing the dynamic interplay between red blood cells (RBCs) and plasma. This study introduces the Core-Plasma Model, a two-phase framework that integrates Newtonian and non-Newtonian elements to represent the RBC-rich core and surrounding CFL. *In vitro* experiments in 25 *µ*m and 50 *µ*m round channels across varying flow rates, hematocrit levels (5–20%), and suspending media (PBS and native plasma) demonstrate the model’s superior ability to capture velocity and shear rate profiles. The Core-Plasma Model offers a robust platform for advancing microscale hemodynamic predictions and deepening the understanding of microvascular flow dynamics.

## Introduction

Blood is a complex suspension of cells in plasma, exhibiting rheological behavior that varies significantly with vessel size, flow conditions, and hematocrit (Ht). In larger vessels, such as arteries and veins, blood behaves approximately as a Newtonian fluid, since the influence of individual red blood cells (RBCs) is averaged out by the dominant plasma flow [1, 21]. However, in the microcirculation—where vessel diameters drop below 100 µm—the discrete nature of RBCs, along with their interactions with plasma and vessel walls, becomes much more pronounced. These microscale effects give rise to non-Newtonian behavior that cannot be ignored in accurate models of microvascular blood flow [22].

One of the most physiologically important phenomena in microvascular blood flow is the formation of the cell-free layer (CFL)—a plasma-rich region near the vessel walls depleted of RBCs. The CFL plays a critical role in reducing flow resistance, improving oxygen delivery, and influencing wall shear stress [1, 15]. Its formation is governed by the interplay of hydrodynamic forces, RBC aggregation, and shear-induced migration [20, 23, 24]. As blood flows through narrow vessels, RBCs tend to migrate toward the center, creating a distinct core-plasma separation.

Blood viscosity, defined as the resistance to deformation under shear, is central to describing flow dynamics. Importantly, blood is shear-thinning—its viscosity decreases with increasing shear rate due to the breakup of RBC aggregates and alignment of cells with the flow direction [2, 6]. This non-Newtonian behavior is further modulated by the Fåhraeus-Lindqvist effect, where apparent viscosity decreases with vessel diameter as the CFL becomes more pronounced [15, 26].

Accurately modeling of blood rheology in the microcirculation is essential for predicting flow behavior and understanding microvascular physiology and pathology. Classical models often fail to fully capture the complexity of blood flow at this scale, particularly under varying hematocrit levels and shear conditions. This study aims to address this gap by systematically evaluating commonly used blood rheology models and proposing an improved framework that explicitly incorporates the impact of CFL formation.

The following section introduces the key rheological and fitting models used to represent blood flow in microvessels. These models range from simple Newtonian approximations to more advanced non-Newtonian formulations, each with varying levels of complexity and physiological relevance.

### Blood Flow Modeling

Several models have been developed to characterize blood rheology in the microcirculation. Each model offers distinct advantages and limitations based on its ability to capture shear-thinning behavior and microstructural effects such as RBC aggregation and CFL formation.

#### Newtonian Model

The Newtonian model assumes constant viscosity and a linear relationship between shear stress and shear rate. The velocity profile in a cylindrical tube is given by:

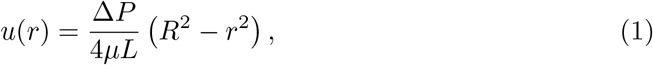

where Δ*P* is the pressure drop, *µ* is fluid viscosity, L is the vessel length, *R* is the vessel radius, and *r* is the radial position from the center of the vessel. Although simple, this model fails to capture the shear-thinning and non-Newtonian effects observed in the microcirculation [2].

#### Power Law Model

The Power Law model introduces non-Newtonian behavior by relating viscosity to shear rate:

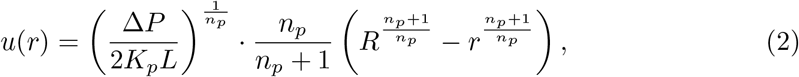

where *K*_*p*_ is the consistency index and *n*_*p*_ is the flow index. This model effectively captures shear-thinning at moderate shear rates, where blood viscosity decreases as shear rate increases. However, it does not account for the viscosity plateau observed at low shear rates, where RBC aggregation increases viscosity, or at high shear rates, where the minimum viscosity is reached. Instead, the viscosity tends to infinity as shear rate approaches zero and to zero at high shear rates—an unrealistic behavior in physiological conditions [2].

#### Carreau Model

The Carreau model addresses the limitations of the Power Law by modeling viscosity over a wide range of shear rates:

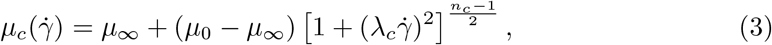

where *µ*_0_ and *µ*_∞_ are viscosities at zero and infinite shear rates, *λ*_*c*_ is the time constant, and *n*_*c*_ is the power index. This model is widely used in hemorheology for its ability to represent the entire shear-thinning spectrum of blood [2]. However, it does not explicitly account for microscale phenomena such as core–plasma phase separation or the formation of a CFL near vessel walls.

### Double-Parameter Power Fit

To improve velocity profiling in microcirculatory flow, a Double-Parameter Power (DPP) Fit was introduced by Koutsiaris et al. (2009) [11]. Unlike traditional rheological models that derive velocity from fluid properties such as viscosity, this approach is a purely empirical fitting model designed to match experimentally observed velocity distributions.

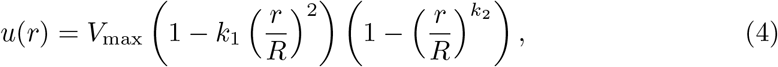

where *k*_1_ controls the velocity gradient near the wall and *k*_2_ controls the bluntness of the core. Although it cannot predict viscosity, it effectively characterizes flow shapes, particularly in the presence of a CFL [10].

### Core-Plasma Model

Building on these foundations, this study introduces a Core-Plasma Model that represents blood flow as a two-phase system: a non-Newtonian, RBC-rich core surrounded by a Newtonian plasma layer near the vessel walls. By explicitly capturing phase separation and presence of a CFL, the model aims to improve predictions of shear-dependent viscosity changes and flow behavior in microvessels, offering greater physiological relevance than traditional single-phase models. The model is validated against experimental microfluidic data collected across varying channel sizes, hematocrits, and suspension mediums. Ultimately, this approach seeks to refine our understanding of microcirculatory blood flow and enhance predictive models for biomedical applications.

## Materials and Methods

### Fluid Preparation

Fresh human blood was collected from a single adult donor under ethics approval from the University of Ottawa (H-03-19-3441). Written informed consent was obtained from the donor prior to participation, as documented using a university-approved consent form. The sample used in this study was collected in March 2024. RBCs were suspended in either phosphate-buffered saline (PBS) or native plasma. Separate samples from the same donor, collected on different days, were used for the 25 *µ*m and 50 *µ*m channel tests, with each channel size tested using a single corresponding sample.

For PBS suspensions, whole blood was centrifuged to remove plasma and buffy coat. The isolated RBCs underwent three PBS washing cycles to eliminate residual plasma proteins, platelets, and white blood cells. RBCs were then re-suspended in PBS supplemented with 0.9 mg/mL glucose to support cell viability and 31.5% OptiPrep^®^ (Sigma Aldrich, Ref. D1556) to stabilize suspension density and minimize sedimentation. Hematocrit (Ht) levels were adjusted to 5%, 10%, 15%, and 20%.

For plasma suspensions, RBCs were similarly washed and re-suspended in density-adjusted native plasma, modified by mixing with dried OptiPrep^®^. This preparation preserved plasma proteins to promote RBC aggregation. Fluorescent tracer particles (0.87 *µ*m, FluoroMax, Thermo Fisher) were added to each sample to enable µPIV measurements.

### Experimental Setup

The experimental system, shown schematically in Fig 1, was designed to simultaneously measure velocity profiles and CFL thickness in microchannels. **Microchannel Fabrication** Microchannels were fabricated using borosilicate glass capillaries (25–50 µm diameter) to model microvessels, following the method developed by Chartrand et al. (2023) [5]. The channels were coated with Poly(L-Lysine)-PEG methyl ether (PLL-PEG) to minimize red blood cell (RBC) adhesion to the glass surface. **Measurement Acquisition** Pressure-driven flow was precisely controlled and monitored using a flow controller and sensor system. Velocity profiles were measured using micro-particle image velocimetry (µPIV) with fluorescent tracers, while CFL thickness was captured in parallel using high-speed imaging. Flow rates were adjusted to produce a physiological yet broad range of shear rates (118–4200 s^−1^), estimated based on the channel dimensions and an assumed blood viscosity of 3.5 cP [2].

**Fig 1.**
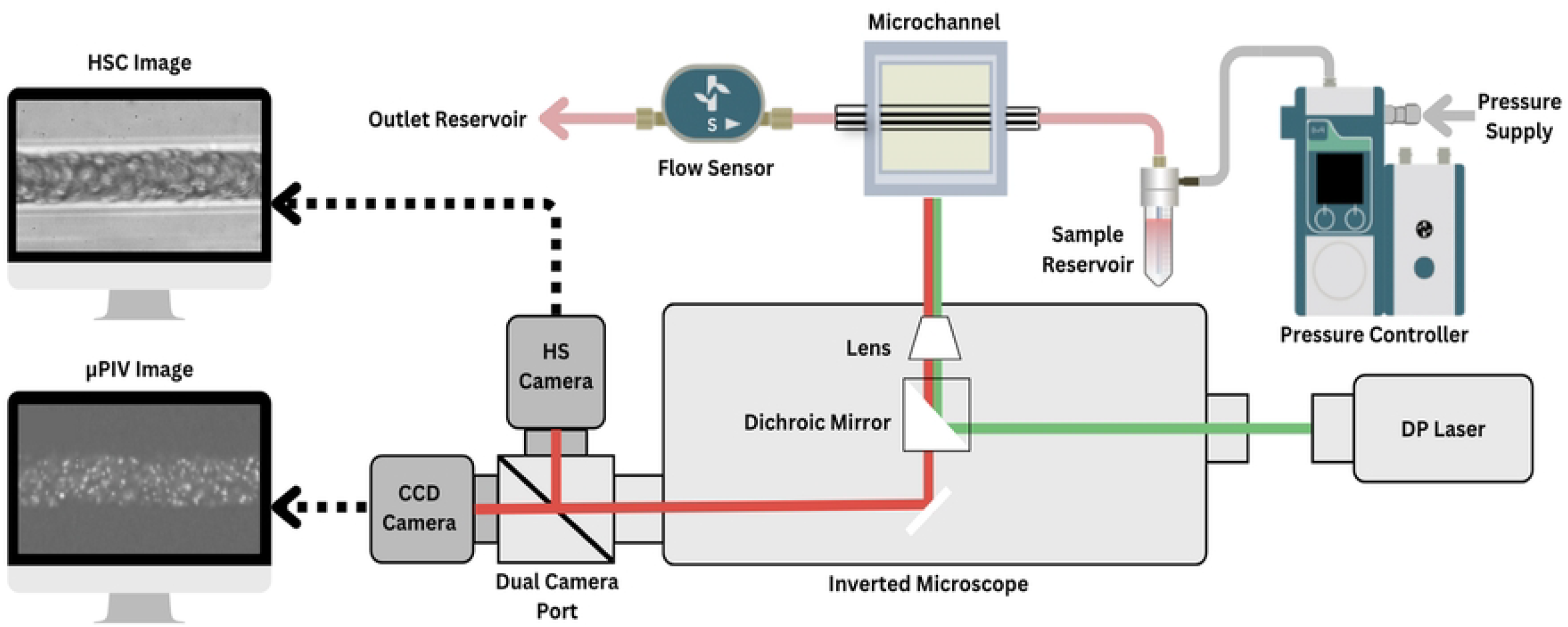
Experimental setup for simultaneous velocity and CFL measurements. Pressure-driven flow was regulated using a Fluigent Flow EZ system, with real-time flow monitoring via an in-line flow sensor. The microchannel, a 25–50 *µ*m diameter borosilicate glass capillary, was mounted on an inverted microscope equipped with a dual-camera port. A 532 nm laser illuminated fluorescent tracer particles for velocity measurements using a LaVision FlowMaster micro-particle image velocimetry (*µ*PIV) system and a CCD camera. Simultaneously, a high-speed camera (HSC) captured red blood cell (RBC) distributions at 100 fps with a 20× objective for cell-free layer (CFL) analysis.

**Image Post-Processing:** For each flow rate, HSC images were collected and imported into a MATLAB application designed for image processing to obtain the optical CFL thickness (*δ*_*o*_), developed by Fenech et al. (2023) [8]. For each flow rate, images were collected and processed using a gradient-based spatiotemporal method to detect the RBC core and compute *δ*_*o*_. For velocity post-processing, image pairs were captured at each flow rate and analyzed using Nguyen’s cross-correlation algorithm with adaptive interrogation windows [17]. Velocity profiles were averaged, and noisy data were filtered based on standard deviation to generate reliable two-dimensional flow fields. The velocity acquisition process is illustrated in Fig 2.

**Fig 2.**
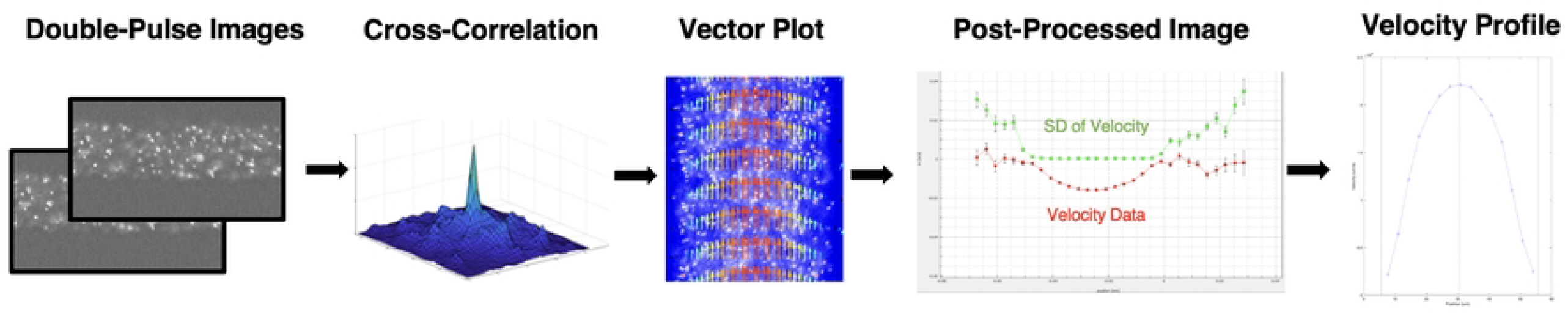
µPIV velocity profile acquisition workflow. Step-by-step process used to extract velocity profiles from microfluidic blood flow experiments using µPIV. (1) Double-pulse images: Sequential images captured using a high-speed camera with fluorescent tracer particles suspended in the flow. (2) Cross-correlation: The Nguyen correlation algorithm [17] is applied to determine the displacement of tracer particles between image pairs. (3) Vector plot: Velocity vectors are generated based on particle displacement, visualizing flow direction and magnitude. (4) Post-processed image: The velocity field is refined by averaging multiple image pairs, with velocity data (red) and standard deviation (green) plotted. Regions where the standard deviation is zero indicate reliable velocity measurements. (5) Velocity profile: Truncated and extracted velocity profile is displayed, showing the parabolic distribution typical of microchannel flow, with a peak at the centerline (red line).

### Mathematical Analysis of Velocity Profiles

#### The Developed Core–Plasma Model

The Core–Plasma Model captures the biphasic nature of blood by dividing the flow domain into two distinct regions: a Newtonian CFL and a non-Newtonian RBC-rich core. This approach enables the integration of both Newtonian and shear-thinning behavior in a single analytical framework.

The governing assumptions include steady, incompressible, laminar, axisymmetric, and fully developed flow in a cylindrical tube, with body forces neglected. The domain is partitioned as follows:

- **Newtonian CFL layer (*R* −*δ*≤ *r* ≤ *R*)**: This region represents the plasma-rich CFL and is modeled as a Newtonian fluid with constant viscosity *µ*_*p*_. The velocity profile is derived by solving the simplified Navier–Stokes equation, applying the no-slip boundary condition at the vessel wall (*r* = *R*) and continuity of shear stress at the core–CFL interface (*r* = *R* − *δ*).
- **Non-Newtonian core (0 ≤ *r < R* − *δ*)**: The RBC-rich core is modeled as a power-law fluid, where viscosity depends on local shear rate. The solution enforces the symmetry condition at the centerline (*r* = 0) and ensures continuity of velocity at the interface (*r* = *R* − *δ*).

This two-region analytical model provides a simplified yet representative approximation of microvascular blood flow, enabling parameter extraction (e.g., flow indices) under physiologically relevant conditions:

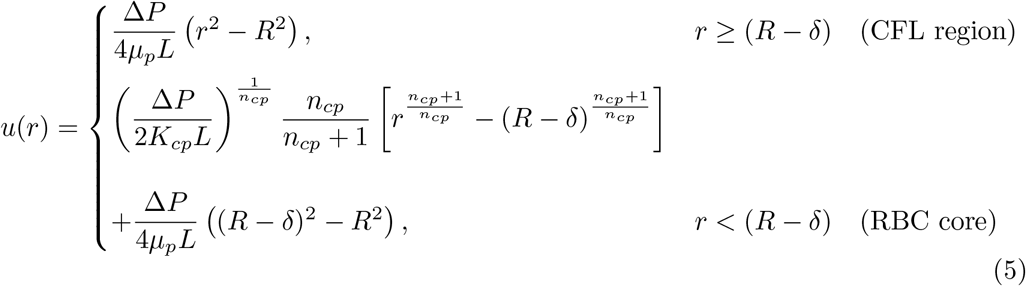

whereδrepresents the CFL thickness.

The shear rate for the Core-Plasma model is given by:

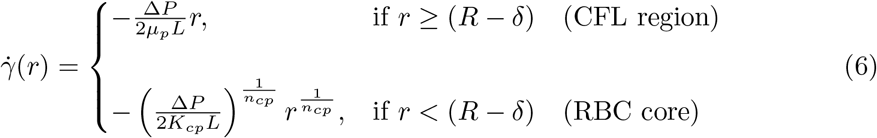

where:

- Δ*P* is the pressure drop,
- *µ*_*p*_ is the Newtonian viscosity of plasma,
- *L* is the vessel length,
- *r* is the radial distance from the channel center,
- *R* is the total vessel radius,
- *δ*is the CFL thickness,
- *K*_*cp*_ and *n*_*cp*_ are the consistency and flow indices governing non-Newtonian behavior.

A velocity distribution, shown in Fig 3, illustrates an example of the Core-Plasma Model overlaid on experimental data, highlighting distinct regions in the RBC-rich core and the surrounding CFL.

**Fig 3.**
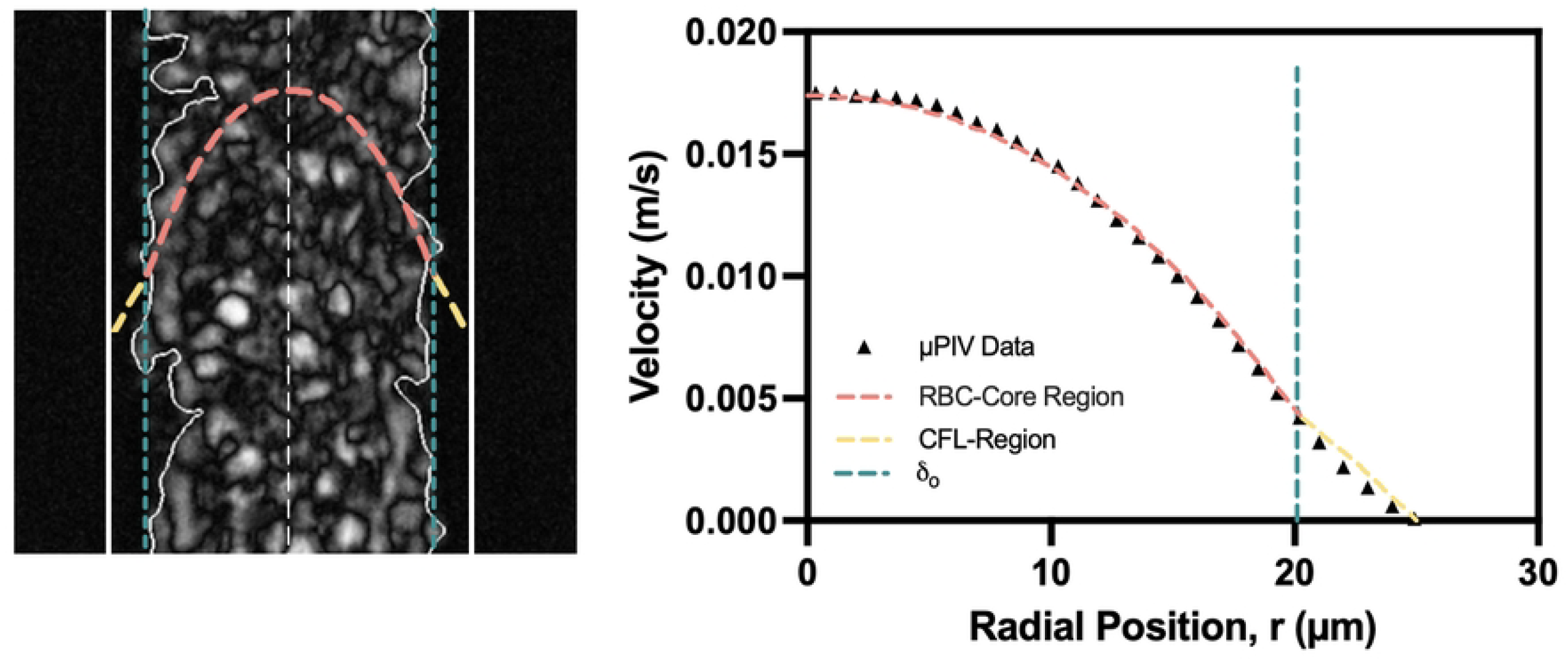
Velocity profile illustrating the Core-Plasma Model in a 50 µm channel. Visualization of the Core-Plasma Model for a 15% hematocrit suspension in plasma flowing through a 50 µm microchannel under 180 mbar pressure. The left panel shows a high-speed image of RBC distribution at high contrast with overlaid core and CFL boundaries. The right panel plots the velocity profile extracted from micro-particle image velocimetry (µPIV) data, with segmented fits for the RBC core (red dashed) and CFL region (yellow dashed). The vertical dashed line indicates the optically measured cell-free layer thickness (*δ*_*o*_).

Direct measurement of plasma viscosity (*µ*_*p*_) is not feasible in our experimental system. Instead, we estimate it using the apparent viscosity (*µ*_*app*_) calculated from Poiseuille’s Law. This estimated value is used as an initial parameter for optimization in the Core-Plasma Model. The approach aligns with the Newtonian assumption applied to the CFL, where plasma is the dominant fluid component.

Poiseuille’s Law assumes laminar flow of a Newtonian fluid with constant viscosity and provides a practical method for estimating *µ*_*app*_ in microcirculatory settings due to its analytical simplicity [16]. In this study, *µ*_*app*_ is computed from experimentally measured flow rate and pressure drop using the expression:

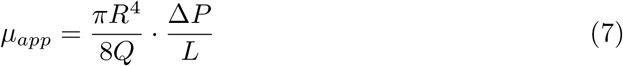

As detailed in the Supporting Information (Fig. 12), the pressure drop across the upstream tubing is negligible, allowing the measured pressure to represent the total pressure gradient across the microchannel. The resulting *µ*_*app*_ serves as an effective estimate for *µ*_*p*_ and provides a baseline for comparing Newtonian and non-Newtonian flow behavior in confined geometries.

### Shear Rate and Hydrodynamic CFL Determination

The shear rate is identified by locating the boundary between the CFL and the RBC core, where an intersection occurs. This intersection is marked by an abrupt change in shear rate 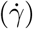, reflecting the distinct rheological properties of the plasma-rich CFL, which behaves more like a Newtonian fluid, and the RBC-dense core, which exhibits non-Newtonian characteristics. The shear rate profile is approximated by two slopes: one originating at the vessel center (*r* = 0) and extending towards an unknown intersection point (ip), and another slope up to the vessel wall (*r* = *R*). The goal is to determine the intersection point, so we can identify the hydrodynamic CFL (*δ*_h_). Mathematically, the shear rate 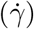 is estimated as:

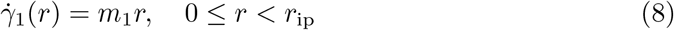

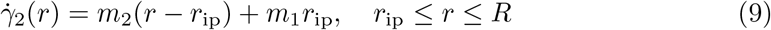

where:

- *m*_1_ is the slope of the first linear region (RBC core),
- *m*_2_ is the slope of the second region (CFL),
- *r*_ip_ is the shear intersection point, representing the hydrodynamic RBC core boundary.

To determine *r*_ip_, we minimize the squared error between the measured shear rate data and the piecewise linear model:

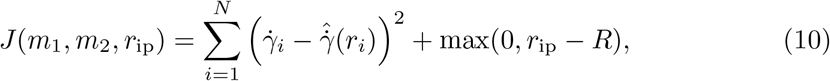

where 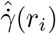 represents the fitted shear rate values and the additional term ensures that the transition point remains within the physical boundary *r*_ip_ ≤ *R*. Once *r*_ip_ is optimized, the hydrodynamic CFL thickness is computed as:

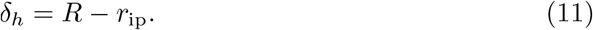

#### Shear Rate at the Intersection

The shear rate at the intersection 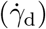 quantifies the rate of deformation at the boundary between the RBC core and the CFL. It is defined as:

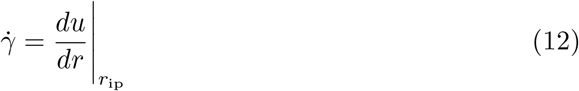

This value represents the final shear rate within the RBC core before transitioning into the CFL, marking the point where the flow dynamics shift from a non-Newtonian to a Newtonian regime. Fig 4 presents a representative shear rate profile, illustrating how linear fits differentiate the RBC core and CFL regions. The plots correspond to half of the microchannel width.

**Fig 4.**
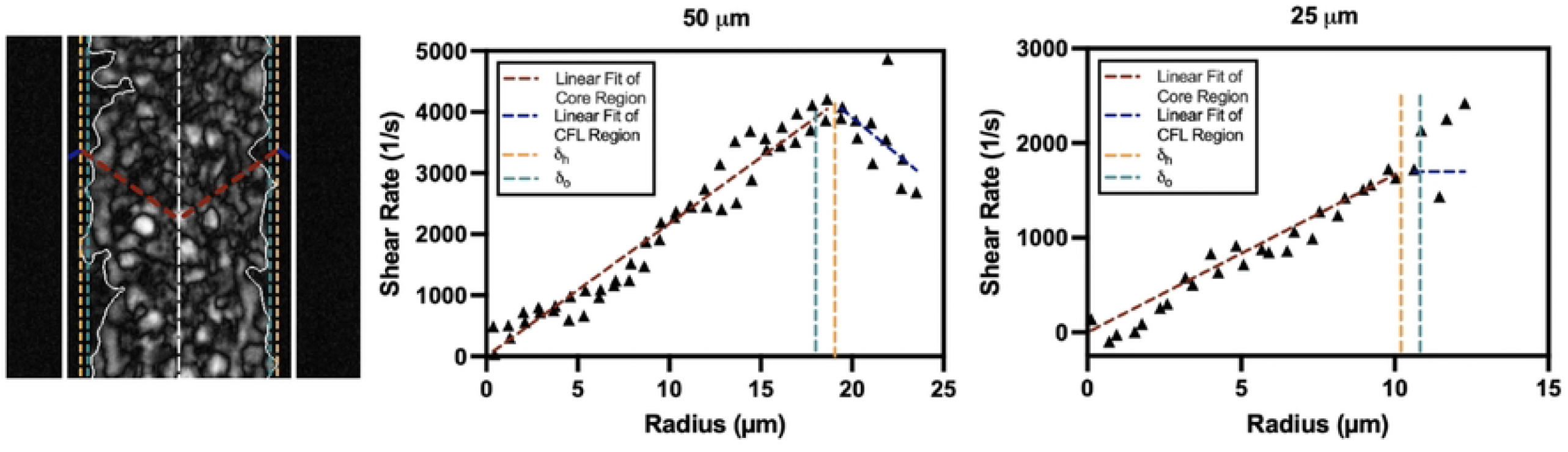
Shear rate at the intersection of a 15% Ht suspension in plasma within 50 µm and 25 µm microchannels. Shear rate profiles for a 15% hematocrit suspension in plasma under an applied pressure of 180 mbar, shown for a 50 µm (left) and 25 µm (right) microchannel. Linear fits are applied to the core and cell-free layer (CFL) regions, with dashed vertical lines indicating the optically measured (*δ*_*o*_) and hydrodynamically estimated (*δ*_*h*_) CFL boundaries. Black triangles represent the numerically derived shear rate obtained from velocity profile data collected via *µ*PIV.

## Results

### Velocity Profiles and Model Performance

Fig 5 and Fig 6 illustrate representative velocity and shear rate distributions for a 15% hematocrit (Ht) suspension in plasma, flowing through 50 µm and 25 µm microchannels under an applied pressure of 180 mbar. The velocity profiles (Fig 5A and Fig 6A) exhibit slightly blunted shapes, with experimental *µ*PIV measurements showing strong agreement with theoretical model fits. In both cases, the boundaries of the CFL, denoted as *δ*_*o*_ and *δ*_*h*_, are well-defined and closely aligned, with a notably thinner CFL observed in the 25 µm channel.

**Fig 5.**
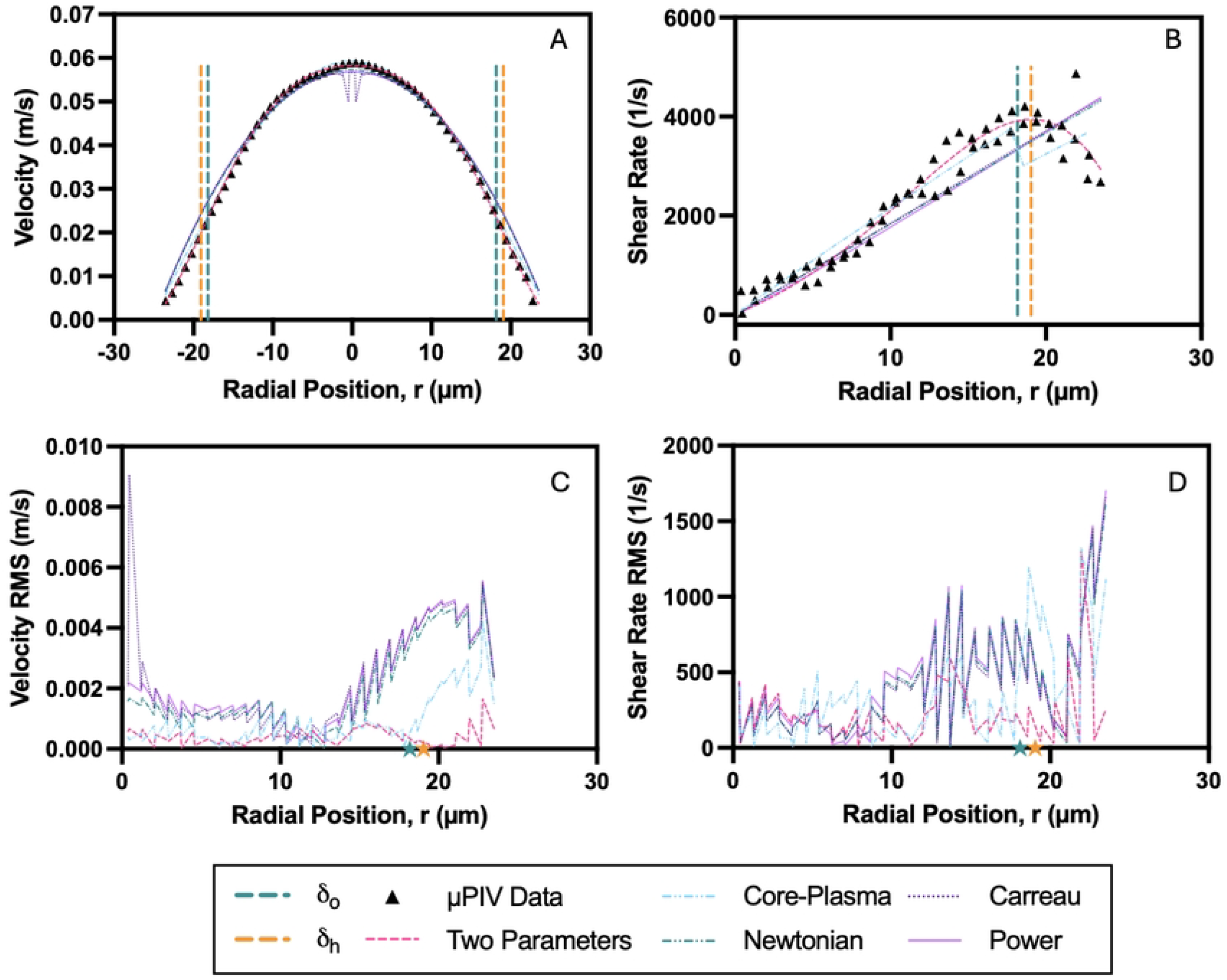
Velocity and shear rate characteristics of a 15% hematocrit suspension in plasma within a 50 µm microchannel. Flow behavior under an applied pressure of 180 mbar is presented: (A) Velocity profile across the channel width, comparing experimental µPIV data (black triangles) with model fits. (B) Shear rate profile derived from the velocity data, with model fits applied to the core and CFL regions. (C) Root mean square (RMS) error of the velocity model fits across the radial position. (D) RMS error of the shear rate model fits across the radial position. Dashed vertical lines or stars represent optically measured (*δ*_*o*_) and hydrodynamically estimated (*δ*_*h*_) CFL boundaries.

**Fig 6.**
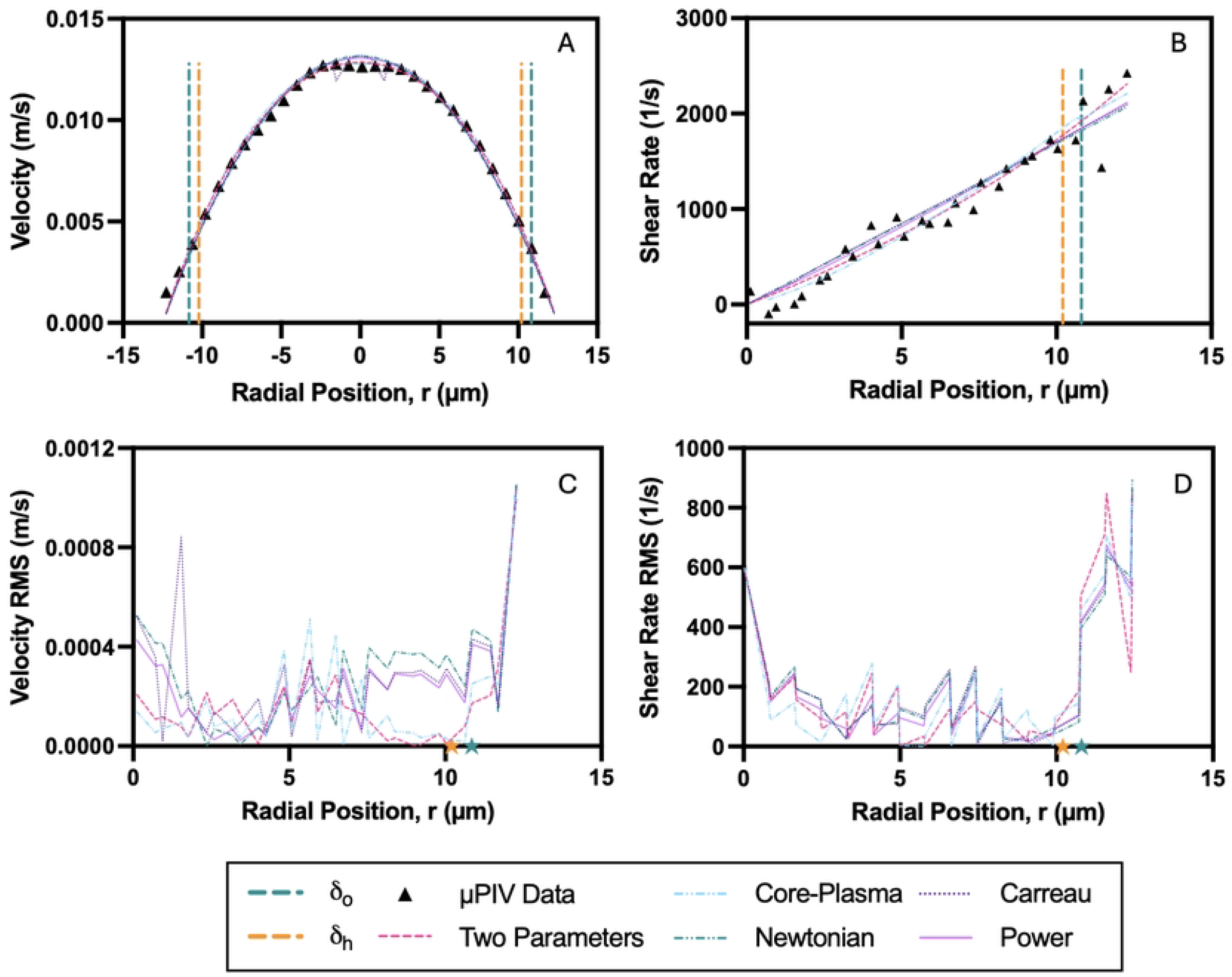
Velocity and shear rate characteristics of a 15% hematocrit suspension in plasma within a 25make µm microchannel. Flow behavior under an applied pressure of 180 mbar is presented: (A) Velocity profile across the channel width, comparing experimental µPIV data (black triangles) with model fits. (B) Shear rate profile derived from the velocity data, with model fits applied to the core and CFL regions. (C) Root mean square (RMS) error of the velocity model fits across the radial position. (D) RMS error of the shear rate model fits across the radial position. Dashed vertical lines or stars represent optically measured (*δ*_*o*_) and hydrodynamically estimated (*δ*_*h*_) CFL boundaries.

Model performance is evaluated using the root mean square (RMS) error between experimental data and fitted profiles, as described in Section S2. Velocity RMS errors (Fig 5C and Fig 6C) remain minimal in the core region but increase near the CFL boundary, reflecting the greater sensitivity of the velocity gradient in these regions. Shear rate distributions (Fig 5B and Fig 6B) reveal a distinct intersection between the RBC-rich core and the surrounding CFL, particularly pronounced in the 50 µm channel.

The RMS errors in shear rate (Fig 5D and Fig 6D) are elevated near the CFL boundary in both channel sizes, with more pronounced discrepancies observed in the 25 µm channel.

A summary of velocity and shear RMS errors is provided in Fig 7. The Core-Plasma model shows the most agreement with experimental data among the rheological models, especially in the 50 µm channel, where phase separation effects such as the formation of a cell-free layer are more pronounced. Here, the Core-Plasma model demonstrates statistically significant reductions in both velocity and shear rate RMS errors compared to all other rheological models (p *<* 0.0001 in most comparisons).

**Fig 7.**
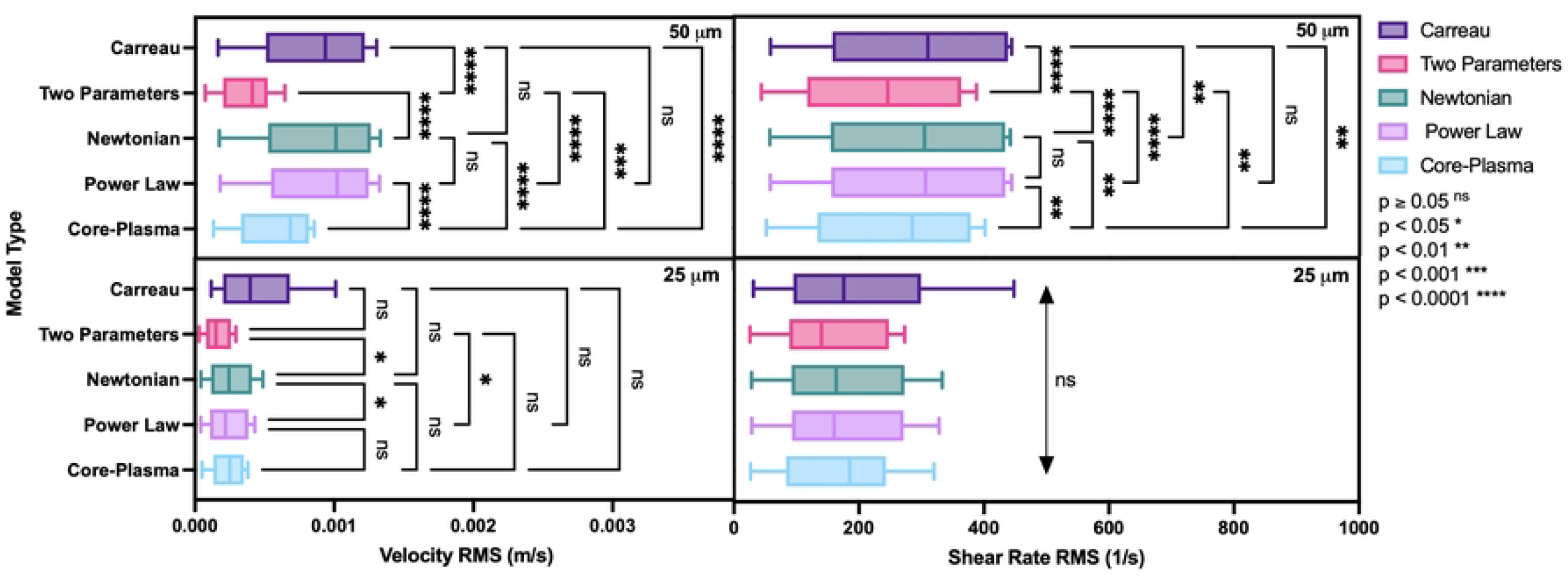
Model comparison of RMS velocity and shear rate errors across microchannel sizes. Root mean square (RMS) values for velocity (left panels) and shear rate (right panels) are compared across five models—Carreau, Double-Parameter Power (Two Parameters), Newtonian, Power Law, and Core-Plasma—using blood suspensions at hematocrit levels of 5%, 10%, 15%, and 20% in both PBS and plasma. Data are grouped by microchannel size: 50 µm (top row) and 25 µm (bottom row). Box plots show the distribution of RMS values across all tested conditions. Statistical comparisons between models were performed using one-way ANOVA with Sidak’s multiple comparisons test. Significance levels are indicated as follows: ns = not significant, * *p <* 0.05, ** *p <* 0.01, *** *p <* 0.001, **** *p <* 0.0001.

The Carreau model consistently exhibits the highest RMS errors, with significant differences observed against all other models in the 50 µm channel for both velocity and shear rate data (p *<* 0.0001), confirming its limitations in accurately describing non-Newtonian blood behavior under confined geometries. The Newtonian and Power Law models yield intermediate performance, showing moderate error levels without significant differences between them in most comparisons, especially in the 25 µm channel where performance across models tends to converge.

Interestingly, model performance diverges more sharply in the 50 µm channel, suggesting that larger channel diameters amplify differences in each model’s ability to capture flow profiles, likely due to enhanced shear gradients and cell-free layer formation. In the 25 µm channel, however, no significant differences were observed between models in shear rate RMS errors, indicating that at small scales, reduced separation between RBC-rich and plasma-rich regions may diminish the influence of complex rheological modeling.

As for the purely empirical fitting, the Double-Parameter Power (DPP) fit shows the lowest RMS velocity errors across most conditions. Statistically, the DPP model performs significantly better than Carreau and Newtonian models in the 25 µm channel (with p *<* 0.05 and p *<* 0.01, respectively), and is not significantly different from Core-Plasma Model.

To further investigate the performance of the Core-Plasma model, we segmented the fitted domain into two distinct regions: the RBC-rich core and the surrounding CFL, and independently evaluated the model’s error in each. As shown in Fig 8, both velocity RMS and shear rate RMS errors were significantly lower in the core region compared to the CFL across both microchannel sizes (50 µm and 25 µm). In the 50 µm channel, the velocity RMS in the core was markedly lower than in the CFL (p *<* 0.0001), and a similar trend was observed for the shear rate RMS (p *<* 0.001). This pattern held true in the more confined 25 µm channel as well, where the core exhibited significantly lower velocity RMS (p *<* 0.001) and shear rate RMS (p *<* 0.01) values than the CFL.

**Fig 8.**
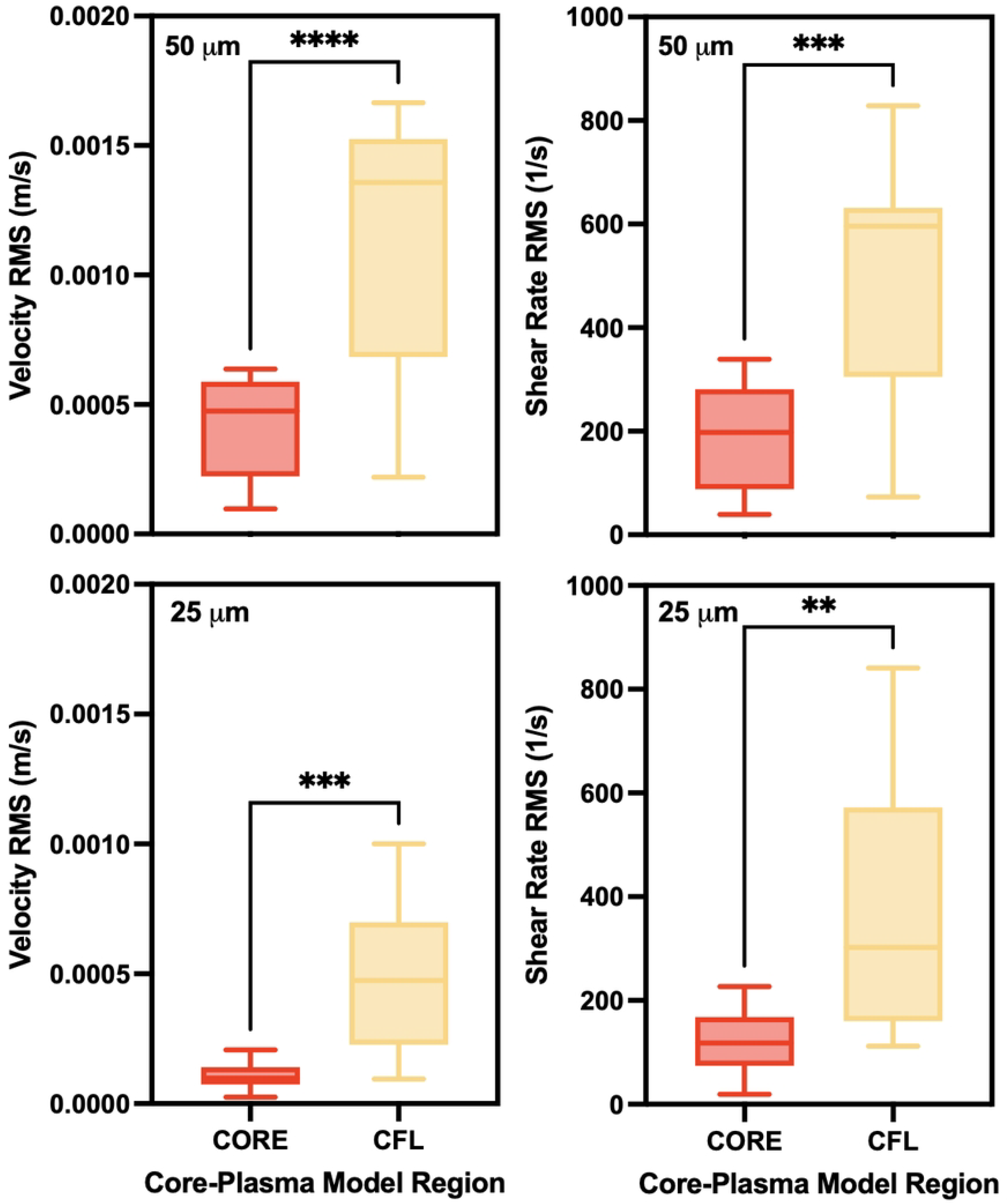
Regional error analysis of the Core-Plasma model in 50 µm and 25 µm microchannels. Velocity (left) and shear rate (right) root mean square (RMS) errors are compared between the core and cell-free layer (CFL) regions of the Core-Plasma model for both 50 µm (top row) and 25 µm (bottom row) channels. Errors were significantly lower in the core region across all cases, indicating better model agreement in the RBC-rich central zone compared to the CFL. Statistical analysis was performed using paired t-tests; significance levels are indicated as follows: \*\**p <* 0.01, \*\*\**p <* 0.001, \*\*\*\**p <* 0.0001.

### Characterization of Non-Newtonian Parameters

Non-Newtonian parameters were extracted by fitting experimental velocity profiles to several rheological models, as summarized in Table 1. Results are reported as mean (standard deviation), with standard deviations shown in parentheses, separately for PBS and plasma suspensions in both 25 µm and 50 µm microchannels, allowing direct comparison of channel size and suspending medium effects. Literature values are included to benchmark physiological relevance and model fidelity.

**Table 1.** Averaged Non-Newtonian parameters by channel size and suspension medium, with literature comparison. Values represent the mean (standard deviation) across all tested flow rates (20–200 mbar) and hematocrit levels (5–20%) for both plasma and PBS suspensions, separated by microchannel size. Individual data points for each condition are provided in the Supplementary Information.

| Model | Parameter (Symbol, Unit) | 25 $\mu\text{m}$ Channel | | 50 $\mu\text{m}$ Channel | | Literature Results |
| --- | --- | --- | --- | --- | --- | --- |
|  |  | PBS | Plasma | PBS | Plasma |  |
| **Experimental-Poiseuille** | Pressure Gradient ($PG$, Pa/m) | $3.21 \times 10^5$ ($1.63 \times 10^5$) | | | | – |
| | Apparent Viscosity ($\mu_{app}$, mPa·s) | 1.98 (0.205) | 1.67 (0.383) | 1.85 (0.309) | 1.74 (0.339) | 1.50 – 2.00 [14] |
| **Carreau Model** | Zero-shear Viscosity ($\mu_0$, mPa·s) | 9.86 (6.17) | 7.27 (5.39) | 13.2 (18.4) | 9.26 (11.8) | 26.9 – 118.6 [6] |
| | Infinite-shear Viscosity ($\mu_\infty$, mPa·s) | 1.86 (0.220) | 1.55 (0.416) | 1.56 (0.356) | 1.75 (0.440) | 1.6 – 2.3 [6] |
| | Time Constant ($\lambda_c$, s) | 3.31 (0.0000931) | 3.31 (0.000322) | 3.31 (0.000477) | 3.31 (0.000576) | 3.312 – 3.313 [6, 12] |
| | Flow Behaviour Index ($n_c$) | 0.354 (0.00902) | 0.382 (0.124) | 0.413 (0.177) | 0.397 (0.155) | 0.353 – 0.369 [6] |
| **Power-Law Model** | Consistency Index ($K_p$, Pa·s$^n$) | 0.00260 (0.000656) | 0.00269 (0.000612) | 0.00297 (0.000435) | 0.00249 (0.000285) | 9.9 – 62.2 [6] |
| | Flow Behaviour Index ($n_p$) | 0.931 (0.0159) | 0.939 (0.0218) | 0.934 (0.0117) | 0.950 (0.0136) | 0.156 – 0.603 [6] |
| **Double-Parameter Power Fit** | Wall Bluntness Shape Factor ($k_1$) | 0.718 (0.469) | 0.374 (0.238) | 0.867 (0.494) | 0.662 (0.269) | 0 – 0.95 [10] |
| | Core Bluntness Shape Factor ($k_2$) | 1.67 (0.429) | 2.53 (2.50) | 3.08 (1.84) | 3.96 (2.56) | 1.74 – 3.66 [10] |
| **Core-Plasma Model** | Consistency Index ($K_{cp}$, Pa·s$^n$) | 0.00310 (0.00519) | 0.00647 (0.0193) | 0.00340 (0.00546) | 0.00263 (0.00388) | – |
| | Flow Behaviour Index ($n_{cp}$) | 1.17 (0.362) | 0.976 (0.278) | 1.02 (0.215) | 1.14 (0.703) | – |

Apparent viscosity, estimated from Poiseuille flow assumptions, was 1.98 (0.21) mPa·s in PBS and 1.67 (0.38) mPa·s in plasma suspensions (25 µm channel), with similar trends observed in 50 µm channels. Plasma-suspended fluid yielded lower viscosity values. All values fall within the physiological range of 1.50–2.00 mPa·s reported by Wells et al. (1962) [14]. The pressure gradient remained consistent across conditions at approximately (3.21 (1.63)) × 10^5^ Pa/m. Fig. 9 illustrates the apparent viscosity’s dependence on pressure gradient, confirming shear-thinning behavior through log–log fitting.

**Fig 9.**
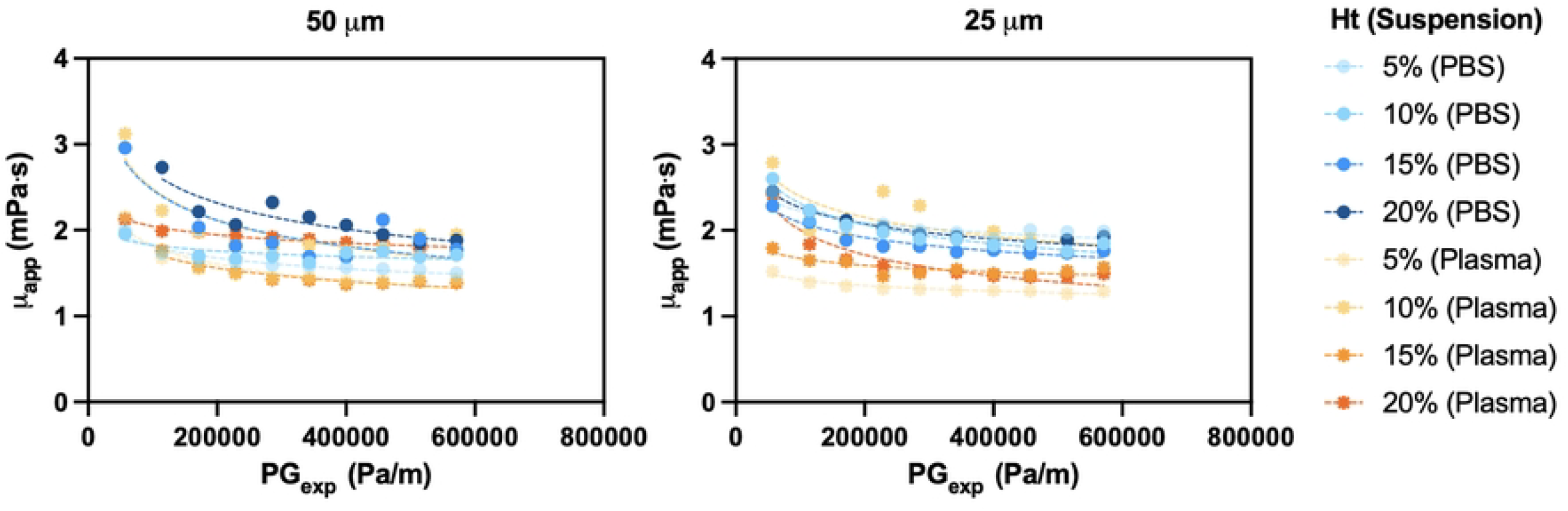
Apparent viscosity as a function of pressure gradient in microchannels. Apparent viscosity, derived from Poiseuille flow assumptions, is plotted as a function of experimental pressure gradient for 50 µm (left) and 25 µm (right) microchannels. Data are shown across different hematocrit levels and suspension types (PBS and plasma). A log–log fitting approach is used to visualize shear-thinning behavior under varying flow conditions.

The Carreau model captured shear-thinning behavior, with infinite-shear viscosities (*µ*_∞_) ranging from 1.55 (0.42) to 1.86 (0.22) mPa·s, closely aligned with previously reported ranges (1.6–2.3 mPa*·*s) [6]. The variation of *µ*_∞_ with experimental pressure gradient across different hematocrits and suspensions is provided in the Supporting Information (Fig. 14), where plasma suspensions have a lower *µ*_∞_, particularly in the 25 *µ*m channel. The time constant (*λ*_*c*_) remained almost identical across all cases (3.31 s with negligible variation). In contrast, zero-shear viscosity (*µ*_0_) showed considerable variability—particularly in PBS suspensions and the narrower 25 µm channel—ranging from 7.27 (5.39) to 13.2 (18.4) mPa·s. These lower values compared to Mehri et al. (26.9–118.6 mPa·s) may reflect differences in hematocrit, channel confinement, and estimation method. This variability of *µ*_0_ with experimental pressure gradient across different hematocrits and suspensions is provided in the Supporting Information (Fig. 13). Fig. 10 displays the Carreau-modeled local viscosity profiles as a function of shear rate, confirming a physiologically realistic shear-thinning response. The 50 µm channel shows a steeper initial drop in viscosity, while the 25 µm channel displays an earlier plateau, indicating reduced shear variation due to geometric confinement.

**Fig 10.**
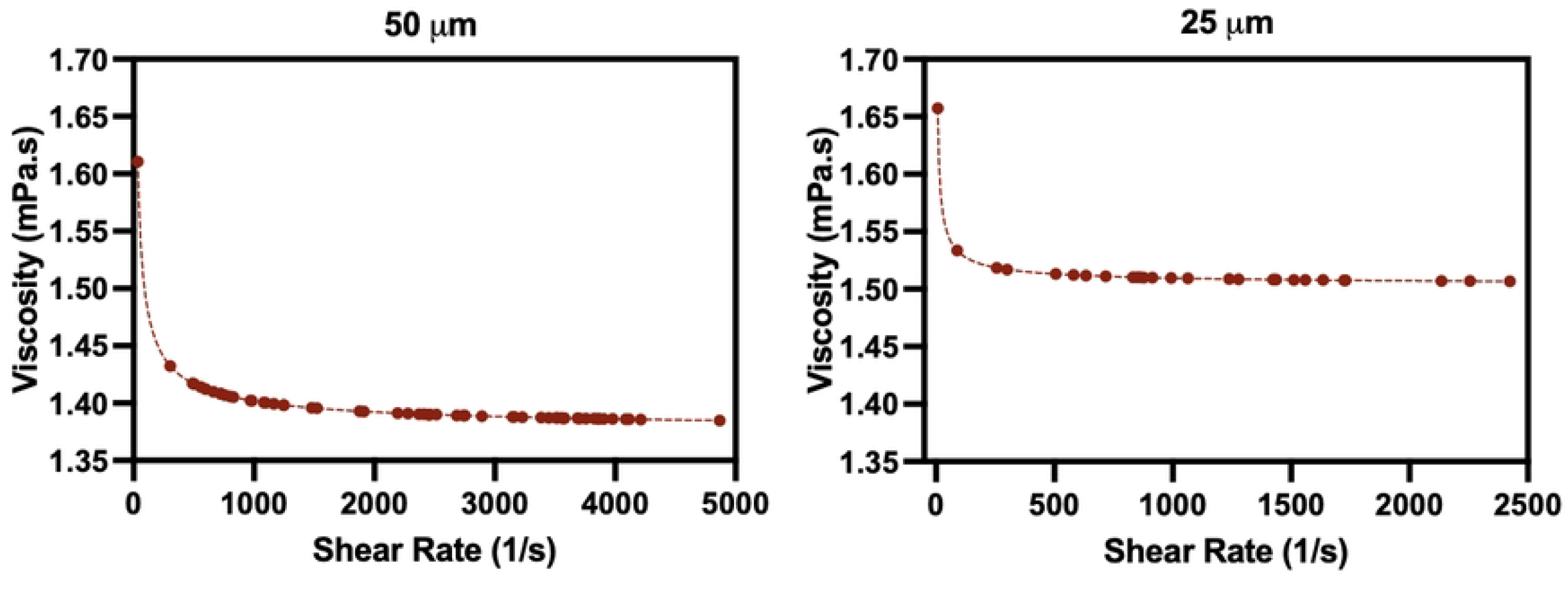
Local viscosity from the Carreau Model as a function of shear rate. Local viscosity is plotted against local shear rate for a 15% hematocrit suspension in plasma flowing through 50 µm (left) and 25 µm (right) microchannels under an applied pressure of 180 mbar. Curves are fitted using the Carreau model with condition-specific rheological parameters to capture shear-thinning behavior.

**Fig 11.**
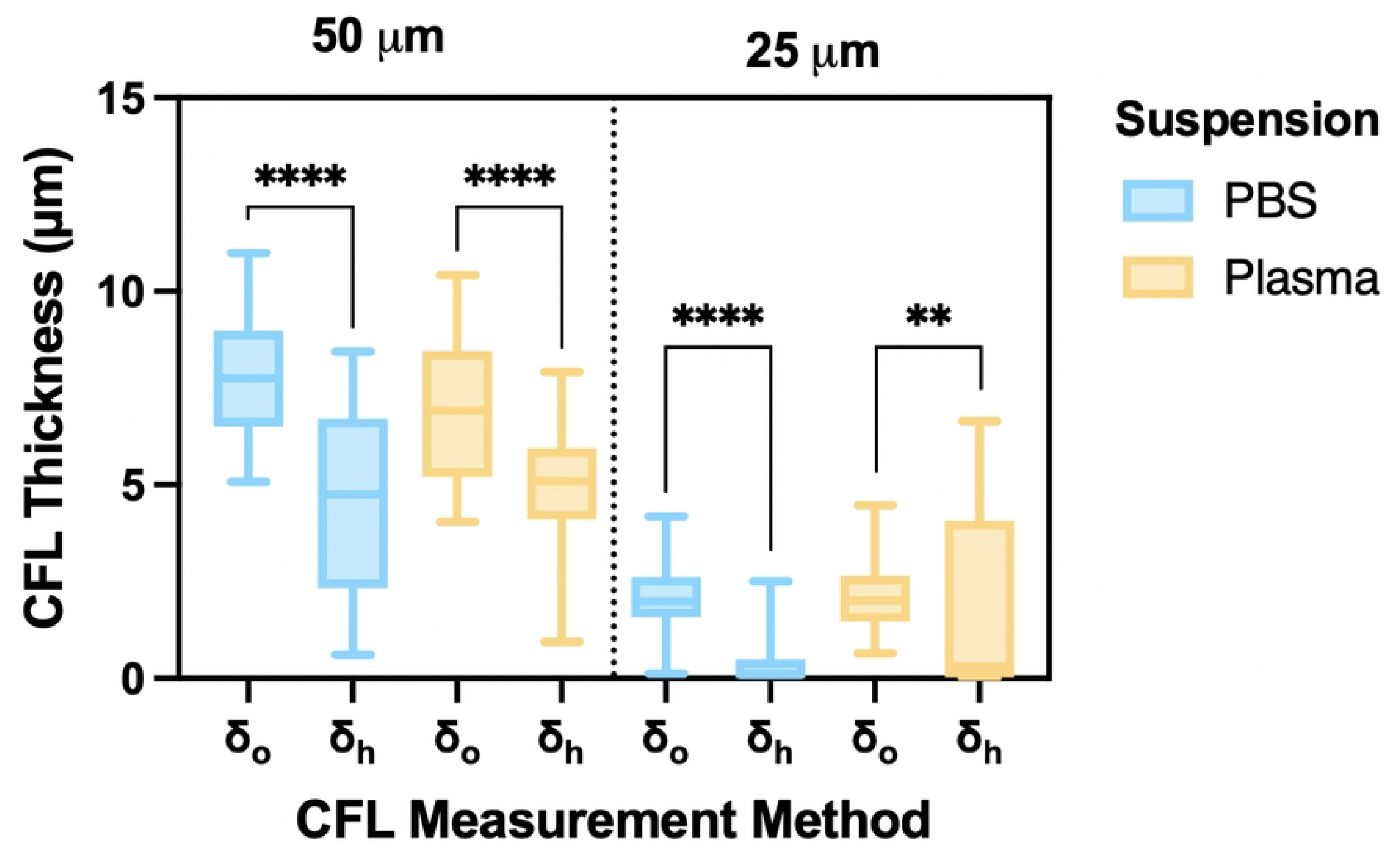
Comparison of cell-free layer thickness across suspensions, channel sizes, and measurement methods. Cell-free layer (CFL) thickness measurements—optical (*δ*_*o*_) and hydrodynamic (*δ*_*h*_)—are shown for both PBS (blue) and plasma (yellow) suspensions at hematocrit levels of 5%, 10%, 15%, and 20% in 50 µm and 25 µm microchannels. Box plots display the distribution of CFL values grouped by measurement method and channel size. Statistical analysis was performed using one-way ANOVA with Sidak’s multiple comparisons test. Significance levels: \*\**p <* 0.01, \*\*\*\**p <* 0.0001.

**Fig 12.**
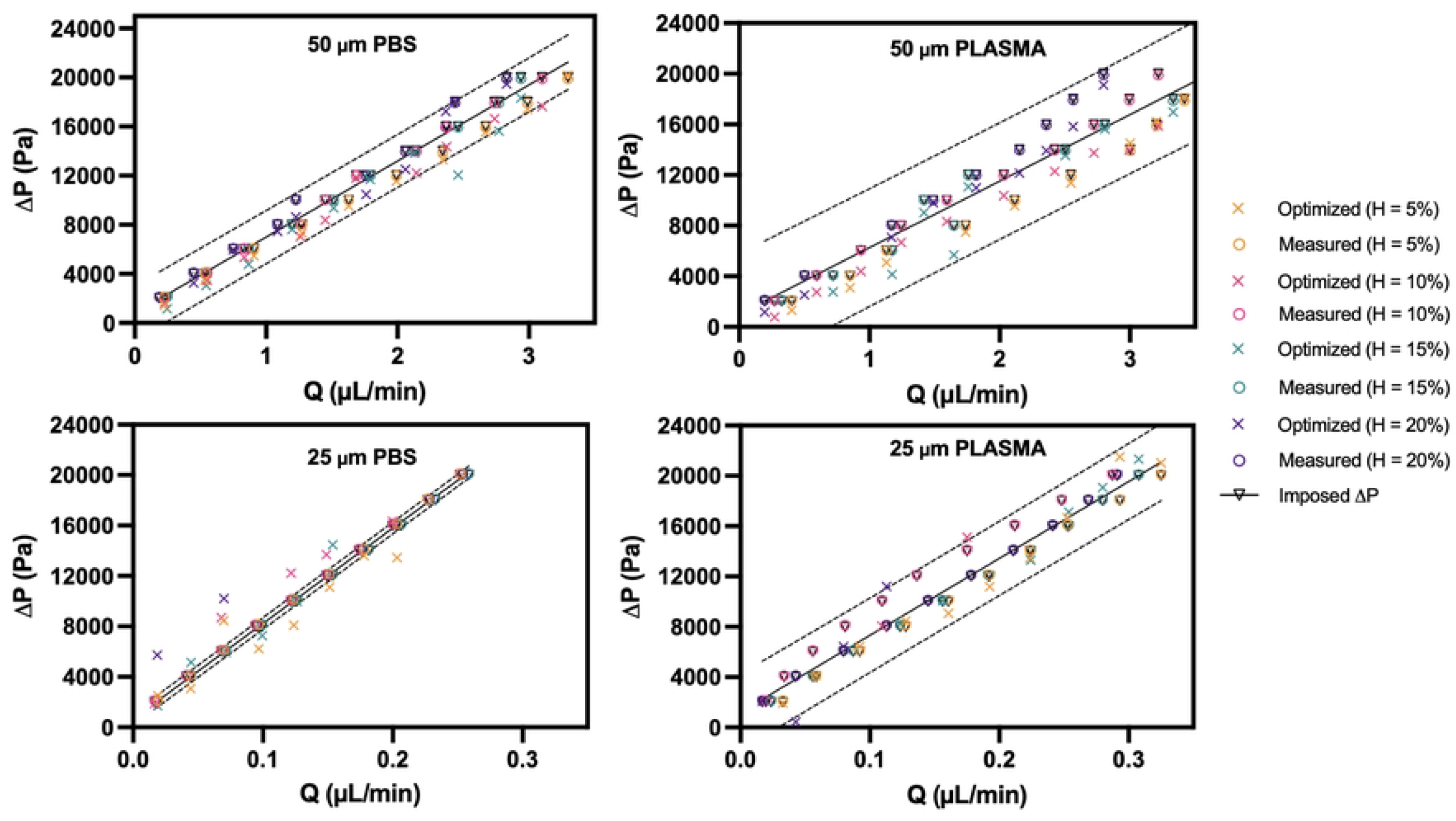
Pressure drop as a function of flow rate in PBS and plasma suspensions. Pressure drop (Δ*P*) is plotted against flow rate (*Q*) for 50 µm (top) and 25 µm (bottom) microchannels, using PBS (left) and plasma (right) suspensions. The **imposed pressure drop** (black triangles) represents the set pressure from the Fluigent system; the **measured pressure drop** (circles) is calculated from direct flow rate data; and the **optimized pressure drop** (crosses) is extracted from model fitting across hematocrit levels (5%, 10%, 15%, 20%). Dashed lines indicate theoretical pressure bounds. Results show strong agreement between imposed and measured values, supporting the validity of imposed pressure as a proxy for flow characterization in confined microchannels.

**Fig 13.**
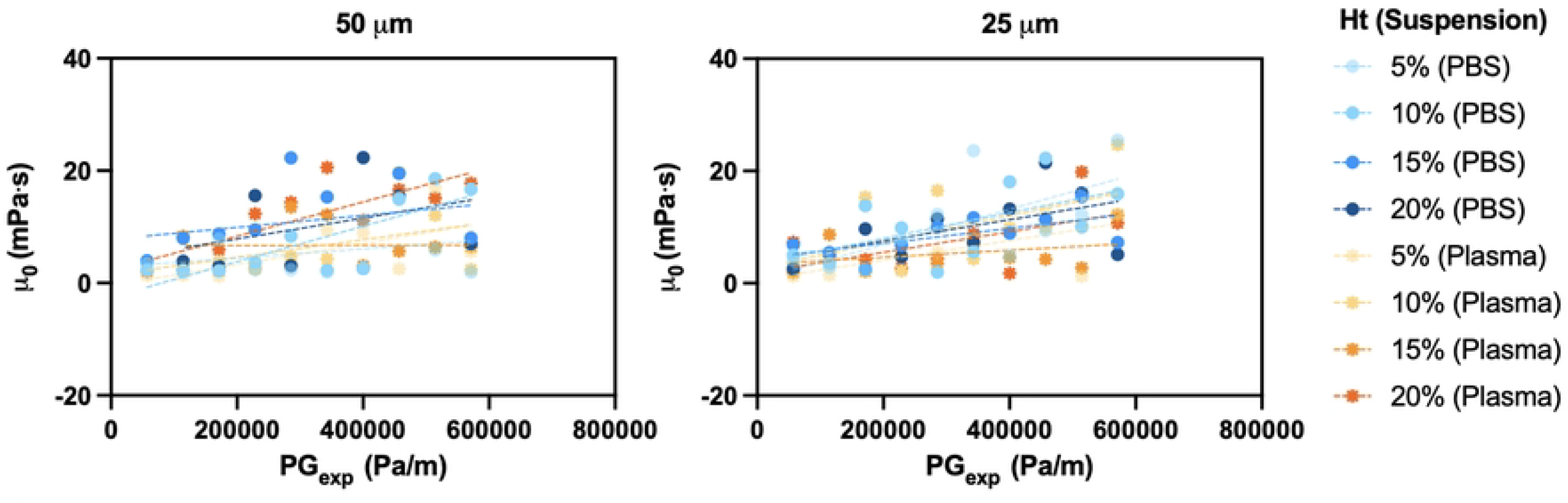
Zero-shear viscosity trends from the Carreau Model across conditions. Zero-shear viscosity (*µ*_0_) derived from the Carreau Model is shown for varying hematocrit levels and suspensions (PBS and plasma) in 50 µm (left) and 25 µm (right) microchannels. Data are plotted as a function of experimental pressure gradient (*PG*_*exp*_) and fitted using simple linear regression to visualize potential trends.

**Fig 14.**
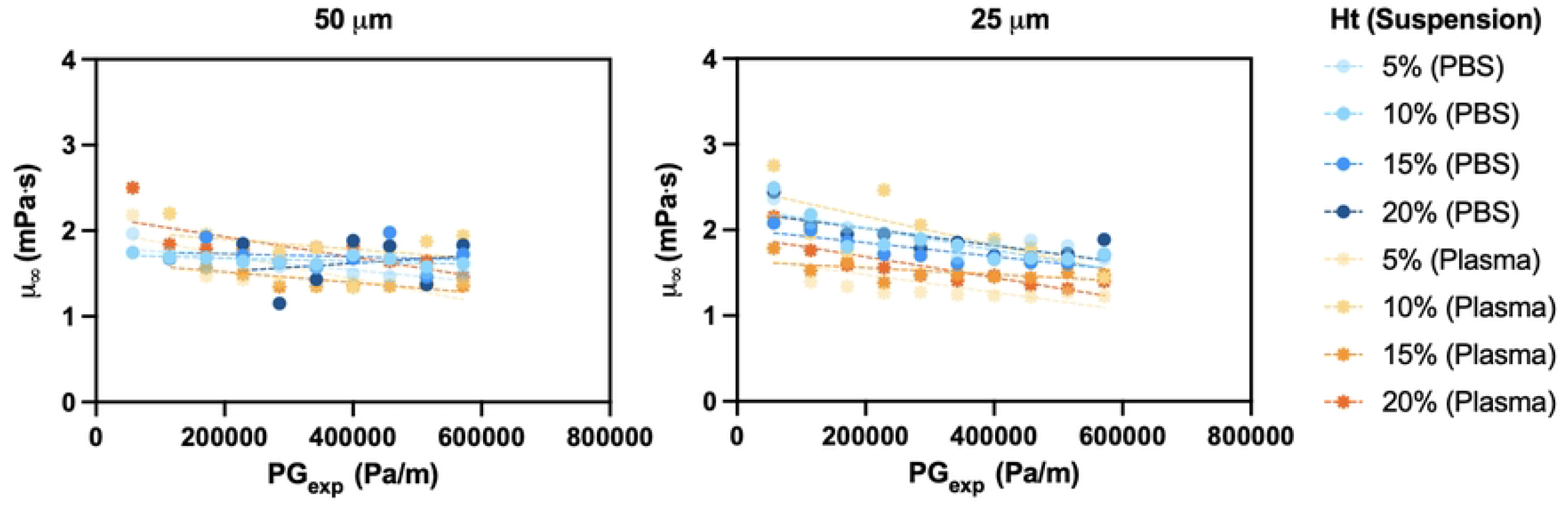
Infinite-shear viscosity trends from the Carreau Model across conditions. Infinite-shear viscosity (*µ*_∞_) predicted by the Carreau Model is shown across varying hematocrit levels and suspension types (PBS and plasma) for 50 µm (left) and 25 µm (right) microchannels. Values are plotted against the experimental pressure gradient (*PG*_*exp*_) and fitted using simple linear regression to assess shear-dependent trends.

The Carreau flow behavior index (*n*_*c*_) ranged from 0.354 (0.009) to 0.413 (0.177), with higher values observed in PBS and larger channels, indicating reduced shear-thinning under these conditions. These values are consistent with the reported physiological range (0.353–0.369) [6].

The Power Law model consistently underestimated non-Newtonian effects, producing flow indices (*n*_*p*_) close to 1 across all conditions (0.931 (0.016) to 0.950 (0.014)), suggesting nearly Newtonian behavior. Its consistency index (*K*_*p*_) was considerably lower than the literature range (9.9–62.2 mPa·s^*n*^), with the lowest values observed in plasma suspensions. The behaviour of *n*_*p*_ and *K*_*p*_ with experimental pressure gradient across different hematocrits and suspensions is provided in the Supporting Information (Fig. 15), where *n*_*p*_ remains near unity and *K*_*p*_ shows no clear trend.

**Fig 15.**
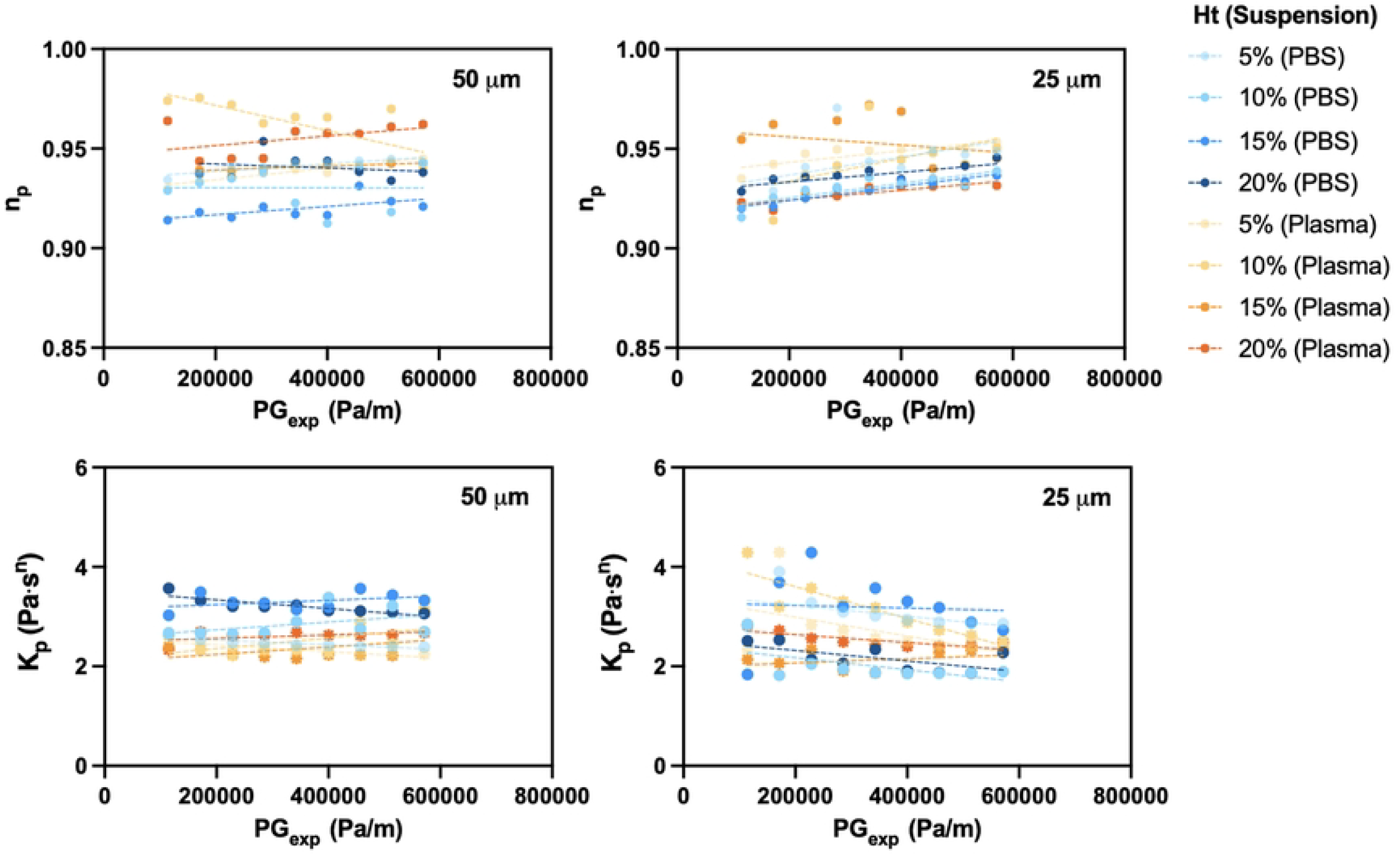
Power Law Model parameters across varying hematocrit levels and channel sizes. Power Law model parameters are shown for different hematocrit levels, suspensions (PBS and plasma), and channel sizes. (Top panel) Flow behavior index (*n*_*p*_) is plotted as a function of experimental pressure gradient (*PG*_*exp*_) for 50 µm and 25 µm microchannels. (Bottom panel) Consistency index (*K*_*p*_) is shown for the same conditions. Simple linear regression is applied to visualize parameter trends across varying flow and confinement conditions.

The Double-Parameter Power (DPP) model effectively captured the shape of velocity profiles, particularly the bluntness at the core and near the walls. Wall shape factors (*k*_1_) ranged from 0.374 (0.238) to 0.867 (0.494), while core shape factors (*k*_2_) reached 3.96 (2.56) in plasma and wide channels. These values fall within the range reported by Pitts et al. (1.74–3.66) [10], confirming the model’s sensitivity to both geometry and fluid phase. The behaviour of *k*_1_ and *k*_2_ with experimental pressure gradient across different hematocrits and suspensions is provided in the Supporting Information (Fig. 16), where *k*_1_ generally increases with *PG*_*exp*_ in narrow channels, while *k*_2_ shows a decreasing trend across most conditions.

**Fig 16.**
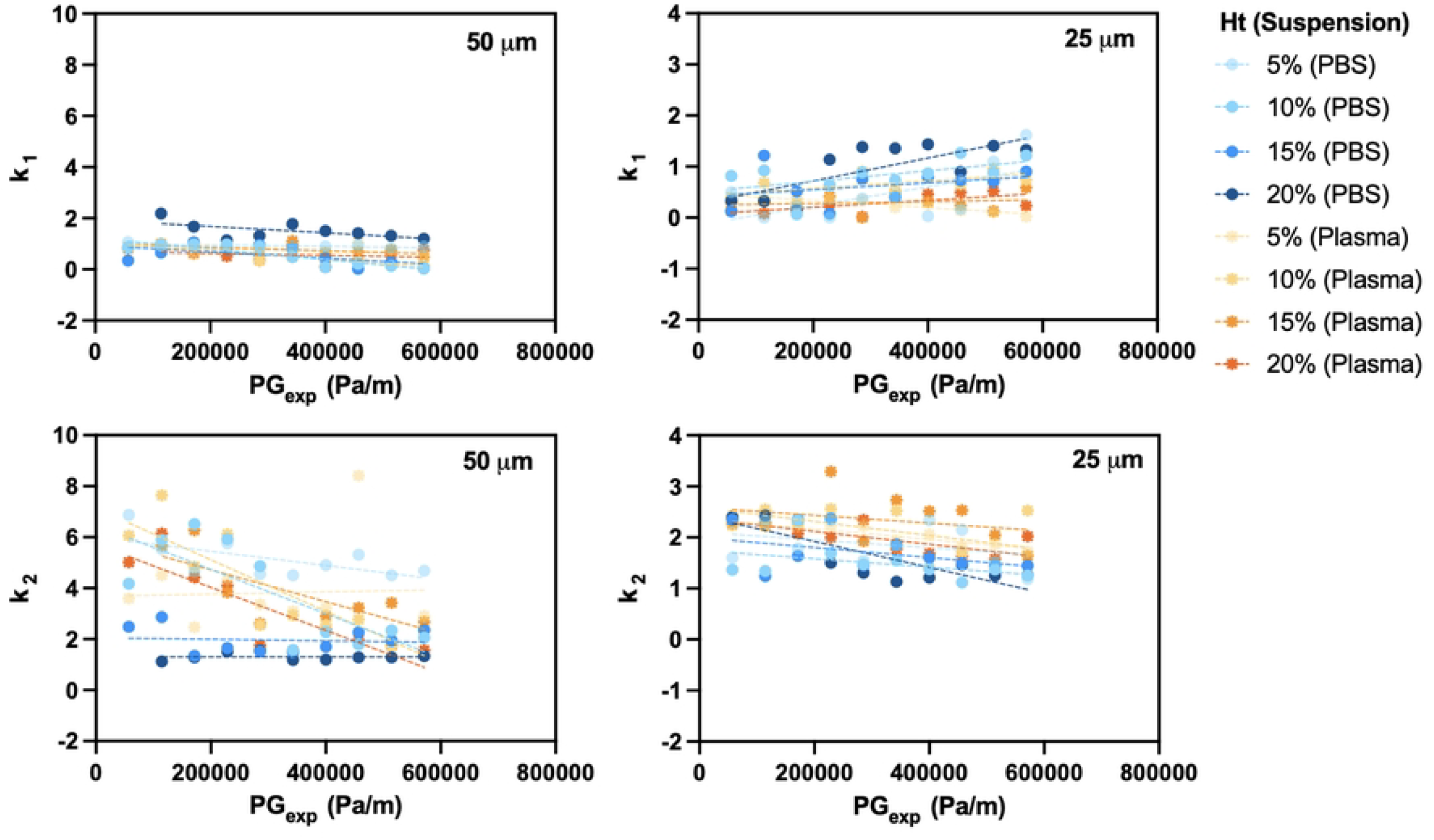
Double-Parameter Power Fit parameters across varying conditions. Variation of the Double-Parameter Power (DPP) Fit parameters—*k*_1_ (core bluntness) and *k*_2_ (wall bluntness)—is shown as a function of experimental pressure gradient (*PG*_*exp*_) for red blood cell (RBC) suspensions in PBS and plasma. Simple linear regression is applied to visualize trends across varying hematocrit levels and channel conditions.

Lastly, the Core-Plasma model produced flow indices (*n*_*cp*_) ranging from 0.976 (0.278) in plasma to 1.17 (0.362) in PBS (25 µm), indicating nearly Newtonian behavior in the RBC-rich core. The consistency index (*K*_*cp*_) was slightly elevated in PBS (0.00310 (0.00519)) compared to plasma (0.00263 (0.00388)), supporting the trend of higher effective viscosity in PBS suspensions. While this model aligns well with observed velocity profiles, the relatively large standard deviations in *K*_*cp*_ values—particularly for plasma—highlights the impact of the CFL that is often neglected in blood modeling. This behaviour of *n*_*cp*_ and *K*_*cp*_ with experimental pressure gradient across different hematocrits and suspensions is provided in the Supporting Information (Fig. 17), where variation is prominent.

**Fig 17.**
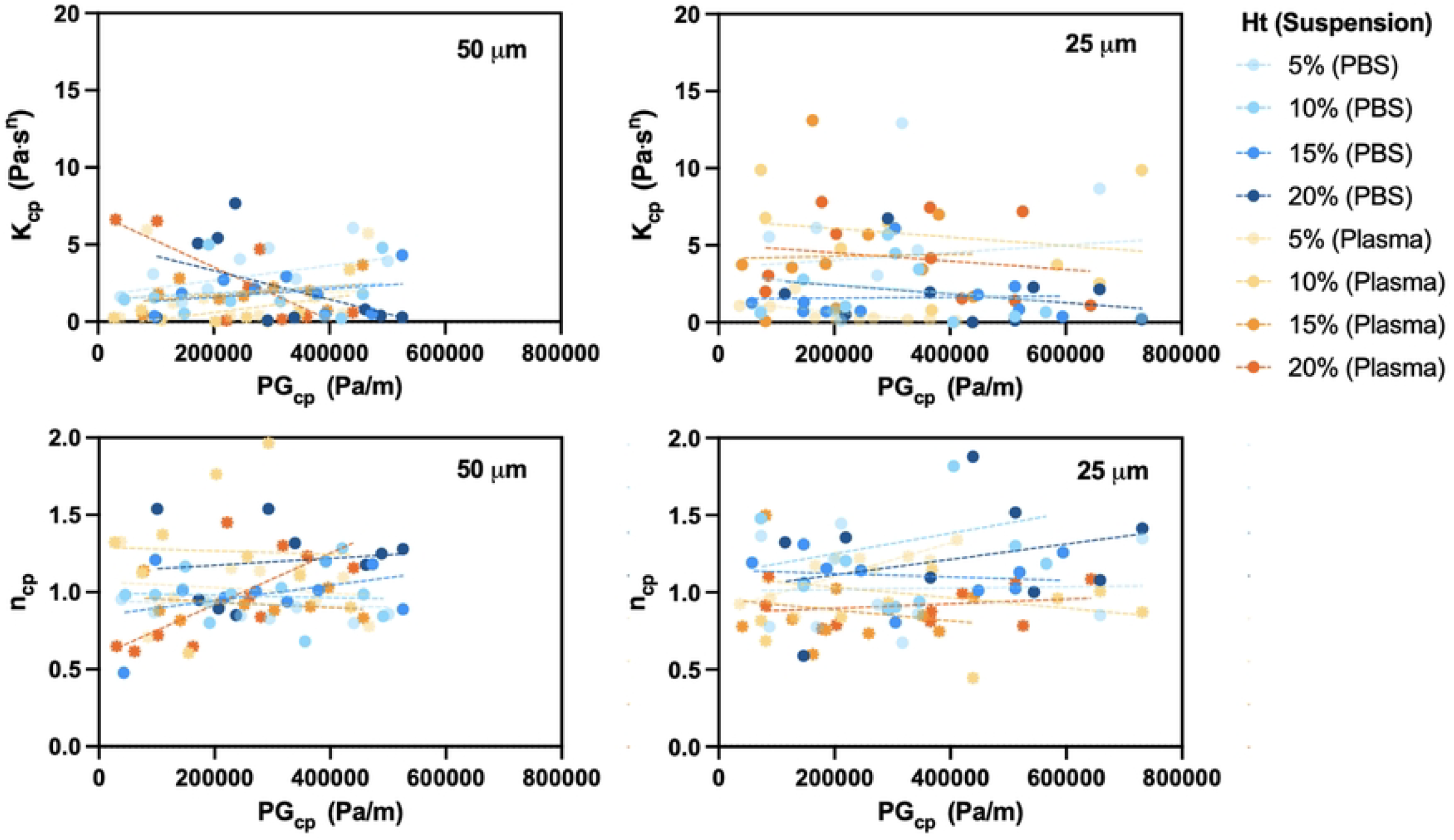
Core-Plasma Model parameters across varying conditions. Trends for the Core-Plasma Model parameters—consistency index (*K*_*cp*_) and flow behavior index (*n*_*cp*_)—are shown as a function of the optimized pressure gradient (*PG*_*cp*_). (Top panel) *K*_*cp*_ values for 50 µm and 25 µm channels. (Bottom panel) *n*_*cp*_ values for the same conditions. Simple linear regression is applied to assess parameter variability, with most cases showing no statistically significant trends.

A detailed summary of the Core-Plasma model parameters across varying pressures, hematocrit levels, and suspension types is provided in Section S3, Table 2. The flow behavior index (*n*_*cp*_) varied between 0.588 and 2.580 across all conditions. For PBS suspensions, *n*_*cp*_ ranged from 0.674 (5% Ht, 160 mbar) to 2.580 (10% Ht, 160 mbar). For plasma suspensions, values ranged from 0.447 (10% Ht, 120 mbar) to 2.094 (10% Ht, 60 mbar). The consistency index (*K*_*cp*_) exhibited a wide distribution. In PBS, values ranged from 9.82 × 10^−5^ (5% Ht, 100 mbar) to 1.29 × 10^−2^ (5% Ht, 160 mbar). In plasma, *K*_*cp*_ ranged from 2.24 × 10^−6^ (10% Ht, 60 mbar) to 1.24 × 10^−1^ (10% Ht, 120 mbar). Across both suspensions, variation in *n*_*cp*_ and *K*_*cp*_ was observed across pressure and hematocrit levels, with no consistent directional trend.

**Table 2.** Core-Plasma model parameters *n*_*cp*_ and *K*_*cp*_ across different pressures, hematocrit levels, and suspensions.

| Core-Plasma $n_{cp}$ | | | | | | | | |
| --- | --- | --- | --- | --- | --- | --- | --- | --- |
| Pressure (mbar) | 5% PBS | 10% PBS | 15% PBS | 20% PBS | 5% Plasma | 10% Plasma | 15% Plasma | 20% Plasma |
| 20 | 1.3661 | 1.4800 | 1.1921 | 1.3244 | 0.9246 | 0.8180 | 0.7786 | 1.0993 |
| 40 | 0.7776 | 1.0433 | 1.3113 | 0.5884 | 0.9641 | 0.6856 | 1.4997 | 0.9147 |
| 60 | 1.2152 | 1.2037 | 1.0603 | 1.3561 | 0.8416 | 2.0940 | 0.8258 | 0.7635 |
| 80 | 0.7764 | 0.8938 | 1.1545 | 0.8888 | 1.1717 | 0.9327 | 0.6000 | 0.7884 |
| 100 | 1.4460 | 0.9056 | 1.1428 | 1.0956 | 1.2265 | 0.8387 | 1.0244 | 0.8120 |
| 120 | 0.9208 | 0.9396 | 0.8065 | 1.8794 | 1.2200 | 0.4465 | 0.7574 | 0.8753 |
| 140 | 0.8604 | 1.3014 | 1.0240 | 1.5176 | 1.1433 | 1.1461 | 0.7343 | 1.0594 |
| 160 | 0.6735 | 2.5798 | 1.0117 | 1.0029 | 1.2331 | 0.9612 | 0.8489 | 0.9910 |
| 180 | 0.8531 | 1.1853 | 1.1308 | 1.0787 | 1.2071 | 1.0077 | 0.7479 | 1.0866 |
| 200 | 1.3496 | 1.8169 | 1.2590 | 1.4137 | 1.3408 | 0.8731 | 0.9712 | 0.7837 |

| Core-Plasma $K_{cp}$ (Pa·s<sup>$n_{cp}$</sup>) | | | | | | | | |
| --- | --- | --- | --- | --- | --- | --- | --- | --- |
| Pressure (mbar) | 5% PBS | 10% PBS | 15% PBS | 20% PBS | 5% Plasma | 10% Plasma | 15% Plasma | 20% Plasma |
| 20 | 7.27E-4 | 6.02E-4 | 1.25E-3 | 1.85E-3 | 1.08E-3 | 9.89E-3 | 3.75E-3 | 3.02E-3 |
| 40 | 5.54E-3 | 2.78E-3 | 7.09E-4 | 3.01E-2 | 1.03E-3 | 6.78E-3 | 6.78E-5 | 1.99E-3 |
| 60 | 7.36E-4 | 1.02E-3 | 1.31E-3 | 5.04E-4 | 2.21E-3 | 2.24E-6 | 3.55E-3 | 7.81E-3 |
| 80 | 6.13E-3 | 5.68E-3 | 6.97E-4 | 6.74E-3 | 2.88E-4 | 6.44E-3 | 1.31E-2 | 5.74E-3 |
| 100 | 9.82E-5 | 4.47E-3 | 7.18E-4 | 1.94E-3 | 2.00E-4 | 4.80E-3 | 8.88E-4 | 7.45E-3 |
| 120 | 3.05E-3 | 3.43E-3 | 6.10E-3 | 1.52E-5 | 1.82E-4 | 1.24E-1 | 3.80E-3 | 4.16E-3 |
| 140 | 4.68E-3 | 3.94E-4 | 2.33E-3 | 1.23E-4 | 2.78E-4 | 7.85E-4 | 5.69E-3 | 1.40E-3 |
| 160 | 1.29E-2 | 6.15E-8 | 1.79E-3 | 2.27E-3 | 1.57E-4 | 3.72E-3 | 3.43E-3 | 1.54E-3 |
| 180 | 8.68E-3 | 6.54E-4 | 8.42E-4 | 2.14E-3 | 1.84E-4 | 2.52E-3 | 7.00E-3 | 1.10E-3 |
| 200 | 3.24E-4 | 7.07E-6 | 3.78E-4 | 1.96E-4 | 6.82E-5 | 9.88E-3 | 1.65E-3 | 7.19E-3 |

### Cell-Free Layer

Fig 11 presents a statistical comparison of CFL thickness measurements, confirming that *δ*_*o*_ is significantly greater than *δ*_*h*_ under all experimental conditions. A one-way ANOVA followed by Sidak’s multiple comparison test is performed, as detailed in Section S2. In the 50 µm channel, the median values of *δ*_*o*_ are 7.89 µm in PBS and 6.91 µm in plasma, while corresponding *δ*_*h*_ values are 4.76 µm and 5.09 µm, respectively. In the 25 µm channel, CFL thickness is markedly reduced, with median *δ*_*o*_ values of 1.12 µm (PBS) and 1.15 µm (plasma), and *δ*_*h*_ values of 0.524 µm and 0.825 µm, respectively.

All comparisons show strong statistical significance (*p <* 0.0001), with the exception of the 25 µm plasma condition, which remains significant but with a lower confidence level (*p* = 0.0065). These findings underscore the influence of both channel size and suspending medium on CFL measurements, with the 50 µm channel exhibiting more pronounced differences between *δ*_*o*_ and *δ*_*h*_ than the 25 µm channel.

## Discussion

### Strengths and Limitations of Rheological Models and Fits

Among the physically grounded models, the Core-Plasma model offers the best overall agreement with experimental data. As shown in Fig 7, the Core-Plasma model consistently yields lower RMS errors than the Newtonian, Power Law, and Carreau models, particularly in the 50 µm channel where phase separation is more pronounced. This strong performance underscores the model’s strength in capturing two-phase flow behavior in microvascular environments. Notably, its ability to account for shear rate intersection between the core and the CFL contributes to its accuracy in reproducing experimental flow fields.

To better understand the model’s internal behavior, we separately analyzed RMS error in the core and CFL regions of the Core-Plasma model (Fig 8). Across both channel sizes, RMS errors were significantly lower in the core region than in the CFL. This pattern reflects the model’s more consistent performance in the RBC-rich zone, where the rheological behavior is relatively stable and dominated by shear-dependent viscosity. Conversely, the higher error observed in the CFL likely arises from increased local variability near the wall, including shear gradients, RBC migration, and potential artifacts in boundary detection. These findings suggest that refining the model’s treatment of the CFL, more accurate plasma viscosity estimation, or improved intersection modeling—could further enhance its predictive power.

Interestingly, the core region error in the 25 µm channel was even lower than that observed in the 50 µm channel, suggesting more RBC packing in highly confined geometries. This increased confinement may lessen lateral migration and flow fluctuations, making the velocity profile within the core more uniform and thus easier to model. However, the overall performance of the Core-Plasma model in the 25 µm channel is slightly diminished due to the reduced CFL thickness and more ambiguous separation between phases, which challenges the two-phase assumptions on which the model is built.

The Power Law and Newtonian models provide intermediate fits. The Power Law model, although better suited than Newtonian assumptions, oversimplifies the system by treating blood as a homogeneous fluid with continuously varying viscosity, thereby ignoring discrete core–CFL dynamics. The Newtonian approximation, assuming constant viscosity, predictably underperforms in shear-thinning-dominated conditions but remains useful as a computationally efficient baseline for benchmarking.

The Carreau model consistently exhibits the poorest fit across both channel sizes. Its limited performance likely results from a mismatch between the model’s target shear range and the experimentally observed shear rates in this study. While the Carreau model is designed to span a wide spectrum of shear rates—from near-zero to extremely high values—the flow regimes probed here are predominantly in the mid-to-high shear range. As a result, the Carreau model fails to appropriately capture the shear-thinning behavior that defines blood flow in confined microchannels under physiological conditions.

The Double-Parameter Power (DPP) fit demonstrates the highest accuracy in capturing velocity profiles across both 25 µm and 50 µm microchannels, as illustrated in Fig 7. It effectively models the characteristic profile bluntness near the centerline, reflecting experimental observations in both RBC-rich and plasma-dominant regions. However, while the DPP model exhibits superior empirical performance, it remains a purely curve-fitting approach lacking a rheological foundation. This absence of physical grounding limits its predictive value under altered physiological conditions or nonstandard geometries, where extrapolation beyond the fitted regime becomes unreliable.

### Interpretation of Non-Newtonian Parameters

The non-Newtonian parameters extracted from experimental velocity profiles reveal consistent and physiologically relevant trends across flow conditions, hematocrit levels, and suspending media. Apparent viscosity, estimated from Poiseuille flow assumptions, remained within the expected physiological range of 1.50–2.00 mPa·s [14]. Plasma suspensions consistently yielded lower *µ*_*app*_ values than PBS across both channel sizes (Fig 9), which is likely due to the enhanced development of the CFL in plasma, reducing near-wall viscosity and overall flow resistance. This effect was especially prominent in the 50 µm channel, where phase separation is more fully developed and pressure gradients are higher.

The Carreau model effectively captured shear-thinning behavior across all conditions. Infinite-shear viscosity (*µ*_∞_) remained consistently between 1.55–1.86 mPa·s, in agreement with values reported by Mehri et al. [6]. The time constant *λ*_*c*_ also showed near-identical values (3.31 s) across all conditions, indicating robust model fitting and consistency. However, the zero-shear viscosity (*µ*_0_) exhibited substantial variability: higher values were consistently observed in PBS (up to 13.2 mPa·s), particularly in the narrower 25 µm channel, while lower values (as low as 7.27 mPa·s) were associated with plasma suspensions. These results suggest that confinement and suspending medium jointly influence RBC aggregation. The absence of plasma proteins in PBS likely promotes stronger hydrodynamic cell–cell interactions and enhanced resistance at low shear, whereas plasma’s dispersive proteins inhibit aggregation, reducing *µ*_0_. Compared to Mehri et al. (who reported values of 26.9–118.6 mPa·s using wider rectangular channels and lower hematocrits), our lower values reflect both our use of cylindrical geometry and our focus on high-shear, physiologically relevant conditions (118–4200 s^−1^ vs. 5–35 s^−1^). The flow behavior index from the Carreau model (*n*_*c*_) increased in PBS and in the 50 µm channel, suggesting that these conditions attenuate shear-thinning and favor more Newtonian behavior. This effect is consistent with the suppression of aggregation at high shear and the more uniform distribution of cells in PBS suspensions. A detailed analysis of the Carreau parameters are outlined in the Supporting Information, Fig 13 and Fig 14.

In contrast, the Power Law model showed limitations in confined microvascular conditions. The flow behavior index (*n*_*p*_) remained between 0.931–0.950 for all conditions, indicating an overestimation of Newtonian behavior and insufficient representation of the non-linear viscosity gradients. Consistency indices (*K*_*p*_) were also notably low, especially in plasma suspensions, and fell well below the range reported by Mehri et al. (9.9–62.2 mPa·s^*n*^). These outcomes highlight the model’s inability to accommodate distinct phase separation and shear-dependent structuring observed experimentally in microchannels. A detailed analysis of the Power Law parameters are outlined in the Supporting Information (Fig. reffig:pparam).

The Double-Parameter Power (DPP) model revealed flow structure behaviour by separating bluntness contributions from the wall and core regions. The wall bluntness parameter (*k*_1_) decreased with increasing pressure in the 50 µm channel, indicating steeper velocity gradients near the wall as RBC aggregation was reduced under higher shear. In contrast, in the 25 µm channel, *k*_1_ increased slightly with pressure, suggesting reduced wall shear gradients potentially due to geometric confinement and limited CFL development. The core bluntness parameter (*k*_2_) exhibited clear pressure-dependent behavior: in the 50 µm channel, *k*_2_ decreased at higher pressure, consistent with more parabolic velocity profiles resulting from RBC disaggregation. However, in the 25 µm channel, *k*_2_ increased under elevated pressure, reflecting a flatter velocity core, likely due to confinement-driven clustering of RBCs. This effect was most pronounced in plasma suspensions, where *k*_2_ reached values as high as 3.96 at low pressure before decreasing toward 1.65 with increasing pressure, indicating a transition from blunt to more parabolic flow. These patterns are consistent with prior findings by Pitts et al. [10] and highlight the DPP model’s sensitivity to confinement, shear rate, and suspension composition. A detailed analysis of the Double-Parameter Power Fit parameters is provided in the Supporting Information (Fig. 16).

The Core-Plasma model demonstrated the highest robustness and physical relevance across conditions. Flow indices (*n*_*cp*_) remained consistent in PBS (1.17 ± 0.36) and plasma (0.98 ± 0.28) in the 25 µm channel, showing with near-Newtonian behavior in the RBC-rich core under high shear rate. Similar trends were observed in the 50 µm channel. As shown in Table 2, values of *n*_*cp*_ were often greater than 1 in PBS, especially at intermediate hematocrits (10%–15%) and pressures between 100–160 mbar. In plasma, *n*_*cp*_ was generally lower, and frequently dropped below 1, indicating stronger shear-thinning and alignment with native plasma behavior. The consistency index (*K*_*cp*_) varied significantly, especially in plasma suspensions. Some plasma conditions yielded extremely low values (e.g., 2.24 × 10^−6^ at 60 mbar, 10% Ht), while others reached peaks above 10^−2^ or even 10^−1^ (e.g., 1.24 × 10^−1^ at 120 mbar, 10% Ht). This large spread reflects spatial heterogeneity within the CFL, likely due to wall effects, RBC margination, or local flow fluctuations not fully captured by the two-phase approximation. By contrast, PBS suspensions exhibited more stable *K*_*cp*_ values across pressures and hematocrit levels, consistent with the absence of protein-mediated aggregation and reduced variability at the wall. No monotonic trends in either *n*_*cp*_ or *K*_*cp*_ were observed with respect to pressure or hematocrit alone. Instead, the results point to an interdependent influence of geometric confinement, suspension composition, and flow rate in shaping the apparent rheology. The increased variability in *K*_*cp*_, particularly in plasma and at intermediate pressures, suggests that improved modeling of the CFL—potentially through spatially resolved viscosity profiles—could further enhance the Core-Plasma framework. A detailed analysis of the Core-Plasma parameters are outlined in the Supporting Information, (Fig. 17). Taken together, these findings highlight the strength of the Core-Plasma model in capturing core behavior and phase separation in microvascular systems, while also pointing to limitations in its treatment of wall-adjacent regions. In future work, region-specific parameterization and direct measurement of local viscosity—particularly within the CFL—may help refine model accuracy for predictive applications in microcirculatory modeling and *in vitro* diagnostics.

### Reliability of CFL Measurements

The comparison between optical (*δ*_*o*_) and hydrodynamic (*δ*_*h*_) measurements of CFL thickness, presented in Fig 11, underscores both the strengths and limitations inherent to each approach. Optical measurements, obtained via high-speed imaging coupled with image processing, provide a more consistent and reliable representation of CFL thickness across experimental conditions. A key advantage of this method is the ability to visually inspect and verify the CFL boundary, offering an additional layer of validation that enhances confidence in the results.

In contrast, the hydrodynamic approach, which derives CFL thickness from *µ*PIV velocity profiles, systematically underestimates the CFL. This discrepancy likely stems from methodological limitations associated with *µ*PIV, including depth-of-correlation effects and out-of-plane motion, both of which compromise the precision of velocity gradient detection near the CFL–core interface [18, 19]. Furthermore, the interface between the plasma-rich layer and the RBC-rich core is not sharply delineated; rather, it constitutes a transitional zone where RBCs intermittently migrate in and out. This dynamic behavior introduces uncertainty into velocity-based measurements, further reducing the reliability of hydrodynamic estimates.

While *µ*PIV remains an invaluable technique for quantifying velocity fields and shear rate distributions, these findings highlight its limitations in accurately resolving sharp spatial transitions such as the CFL boundary. For applications requiring precise measurement of CFL thickness in microvascular flow studies, optical methods should be prioritized due to their higher spatial resolution and reduced susceptibility to artifacts.

## Conclusion

Among the rheological models evaluated, the Core-Plasma model exhibited the strongest predictive capability, effectively capturing the two-phase characteristics of microvascular blood flow, including the separation between the red blood cell (RBC)-rich core and the plasma-rich cell-free layer (CFL). Its ability to reproduce shear rate discontinuities near the channel wall, particularly in the 50 µm geometry where phase separation is more pronounced, reinforces the relevance of two-phase modeling in confined microcirculatory environments.

Despite its strengths, the Core-Plasma model is limited by simplifying assumptions—most notably, the approximation of plasma viscosity using apparent viscosity values and the use of linear fitting approaches that may overlook localized velocity gradient complexities. Additionally, discrepancies between hydrodynamic CFL thickness (*δ*_*h*_), derived from *µ*PIV velocity profiles, and optical CFL thickness (*δ*_*o*_), derived from high-speed imaging, emphasize the need for caution when interpreting near-wall flow behavior based on velocity data alone.

The Double-Parameter Power (DPP) fit, while purely empirical, consistently provided the most accurate fits across all conditions, effectively capturing velocity profile bluntness and wall gradients. However, its lack of physical interpretability limits its utility for extrapolative or mechanistic modeling.

Future work should prioritize direct measurement of plasma viscosity, improved spatial resolution near CFL boundaries, and integration of more physiologically grounded parameters into model formulations. These enhancements would improve model accuracy, strengthen physiological relevance, and support applications in microfluidic design and investigation of pathological microvascular disorders.

## Supporting information

**S1. Comparison of Imposed, Measured, and Optimized Pressure Drops Across Different Hematocrits and Channel Sizes**. The imposed pressure gradient (∇*P*) was used as the driving force in the microfluidic setup to ensure controlled and reproducible flow. To validate its accuracy, we compared the imposed Δ*P* to measured and optimized values. Linear regression models were fitted for each dataset, yielding the following equations: *Y* = 6200*X* + 794.5 (*R*^2^ = 0.9823) for 50 *µm* PBS, *Y* = 5256*X* + 1066 (*R*^2^ = 0.9196) for 50 *µm* plasma, *Y* = 75685*X* + 662.4 (*R*^2^ = 0.9992) for 25 *µm* PBS, and *Y* = 61129*X* + 1209 (*R*^2^ = 0.9670) for 25 *µm* plasma. The high *R*^2^ values confirm the strong linear relationship between flow rate and pressure drop for the 25 *µm* channels, indicating minimal deviation from theoretical expectations. The slight deviations in the plasma cases are expected due to hematocrit-dependent viscosity variations. To determine the optimized Δ*P*, we used an unconstrained fitting approach that adjusted viscosity and shear rate parameters to best match experimental flow characteristics. The optimized Δ*P* values were then compared against the imposed and measured values. Measured Δ*P* was obtained by flowing water through the tubing system without the microfluidic chip to capture intrinsic resistance from the setup. The imposed Δ*P* closely aligned with the optimized and measured values, confirming its reliability for imposing this value as an initial condition in the code. This suggests that, for these experimental conditions, the pressure drop in the tubing can be considered negligible and does not significantly influence the overall flow behavior. However, slight deviations observed in plasma cases indicate that non-Newtonian effects and hematocrit-dependent variations may introduce minor discrepancies, which should be considered when working with higher-viscosity suspensions.

**S2. Statistical Analysis Methods**. Statistical analyses and data visualization were performed using GraphPad Prism (GraphPad Software, San Diego, CA, USA). The analyses included Root Mean Square (RMS) error calculations, ordinary one-way analysis of variance (ANOVA), simple linear regression, and significance testing with a confidence level of 95%.

### Root Mean Square (RMS) Error

The Root Mean Square (RMS) error quantifies the deviation between experimental data and model predictions, providing a measure of model accuracy. It is calculated as:

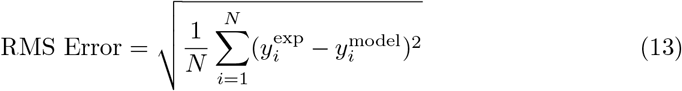

where 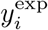 represents the experimentally measured values, 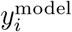 are the corresponding theoretical predictions, and *N* is the total number of data points. A lower RMS error indicates a better fit between the model and experimental data, while a higher RMS error suggests greater discrepancies.

### Ordinary One-Way ANOVA with Sidak Multiple Comparisons

Ordinary one-way Analysis of Variance (ANOVA) is used to determine whether there were statistically significant differences between multiple experimental groups. ANOVA tests the null hypothesis that all group means are equal by comparing the variance within groups to the variance between groups, yielding a p-value that indicates the probability of observing the data if the null hypothesis were true. If a significant difference was detected (p *<* 0.05), Sidak’s multiple comparison test was applied as a post hoc analysis to identify which specific groups differed from each other.

The significance levels are denoted as:

- **p** *<* **0.05** (*) – Significant difference
- **p** *<* **0.01** (**) – Strong significant difference
- **p** *<* **0.001** (***) – Highly significant difference
- **p** *<* **0.0001** (****) – Extremely significant difference

**S3. Core-Plasma model parameters** *n*_*cp*_ **and** *K*_*cp*_ **across different pressures, hematocrit levels, and suspensions**. This section presents the fitted Core-Plasma model parameters *n*_*cp*_ (flow behavior index) and *K*_*cp*_ (consistency index) for RBC suspensions under varying inlet pressures, hematocrit levels, and suspending media (PBS or plasma). These values are obtained by fitting experimental velocity profiles to the Core-Plasma model, enabling quantitative comparisons of non-Newtonian behavior across conditions (Table 2).

**S4. Characterization of Non-Newtonian and Fitting Parameters**. This section presents supplementary figures related to the characterization of blood flow behavior in microfluidic channels.Model-derived parameters are shown across varying hematocrit levels, suspending media (PBS and plasma), and channel diameters (25 µm and 50 µm). Analyses include curve fitting using three non-Newtonian rheological models—Carreau (Fig. 13, Fig. 14), Power Law (Fig. 15), and Core-Plasma (Fig. —as well as the Double-Parameter Power (DPP) Fit (Fig. 16), which is used to capture geometric features of the velocity profile (core and wall bluntness).

## Acknowledgments

The authors would like to thank Andy Vinh Le for his valuable assistance with data interpretation and software implementation, and Camille Chartrand for providing training and guidance on microchannel fabrication.

## Author Contributions

**Conceptualization:** Maya Salame.

**Data curation:** Maya Salame.

**Formal analysis:** Maya Salame.

**Funding acquisition:** Marianne Fenech.

**Investigation:** Maya Salame.

**Methodology:** Maya Salame, Marianne Fenech.

**Project administration:** Marianne Fenech.

**Resources:** Marianne Fenech.

**Software:** Maya Salame, Marianne Fenech.

**Supervision:** Marianne Fenech.

**Validation:** Maya Salame.

**Writing – original draft:** Maya Salame.

**Writing – review & editing:** Marianne Fenech.

## Notes

### Competing Interest Statement

The authors have declared no competing interest.

